# Genetic diversity, predictive protein structures, and interaction networks of Cysteine-Rich Receptor-Like Kinases in *Arabidopsis thaliana*

**DOI:** 10.1101/2025.11.26.690834

**Authors:** Jente Stouthamer, Danilo Pereira, Sergio Martin-Ramirez, Sumanth Mutte, G. Adam Mott, Elwira Smakowska-Luzan

## Abstract

Cysteine-rich receptor-like kinases (CRKs) are a large subfamily of plant receptor-like kinases (RLKs) implicated in immunity and development, yet their ligands, interaction partners, and mechanistic roles remain poorly defined. We combined population-genetic analyses and AlphaFold-based structural prediction to characterise the Arabidopsis thaliana CRK family. Phylogenetic reconstruction from 69 natural accessions resolved five well-supported CRK clades. Nucleotide diversity (π) and neutrality tests revealed heterogeneous diversity across loci, with evidence of both positive and negative selection pressure acting on different CRKs. AlphaFold models of CRK extracellular domains (ECDs) recapitulate the DUF26 structure observed in Plasmodesmata Localizing Protein (PDLP)5/PDLP8 and ginkbilobin-2 but display distinct biochemical properties and disulfide-bond topologies. Pairwise AlphaFold dimer modelling of all 780 CRK-ECD combinations produced 145 high-confidence interaction models; ∼78% of these adopt a shared dimer conformation characterized by an extended intermolecular β-sheet at the interface. Integrating evolutionary and structural approaches reveals clade-specific selective regimes and conserved structural features of CRK ECDs that likely underpin receptor–receptor interactions. Predicted high-confidence dimer interfaces suggest a general mode of CRK-ECD association that can guide targeted biochemical and genetic validation, accelerating functional dissection of this important receptor family.

## Introduction

Plant cells continuously perceive and respond to fluctuating environmental cues via extensive receptor-mediated signalling networks. *Arabidopsis thaliana* encodes >600 receptor-like kinases (RLKs), among which the Cysteine-Rich Receptor-like Kinases (CRKs; 44 members in Arabidopsis) constitute a major RLK subfamily implicated in immunity, stress response and development^1^. Like canonical receptors, CRKs are also modular membrane proteins. At the N-terminus, CRKs have a signal peptide that directs the protein to the plasma membrane. Following that is the extracellular domain (ECD) containing two tandem domains of unknown function (DUF26-A and DUF26-B) defined by a conserved cysteine motif (C-8X-C-2X-C)^2,3^. Canonically, as demonstrated for other RLKs, the ECDs are most likely required for the perception of signalling molecules and complex formation with other RLKs^4^. A long juxtamembrane region (average ∼35 AA) rich in proline residues connects the ECD to the single transmembrane domain (TM) composed of α-helix. Intracellularly, another juxtamembrane region rich in lysine and arginine residues connects the TM to the serine/threonine kinase domain (KD). In general, both the juxtamembrane regions and the TMD can regulate protein activity, localisation, and interaction at the plasma membrane. The conserved KD is required for intracellular interactions and signal propagation through autophosphorylation and the transphosphorylation of target proteins^5,6^. The KD is highly conserved among CRKs, including regions important for function and regulation: the ATP-binding site, catalytic loop, and activation loop.

Functional specificity among RLKs is generally conferred by ECD architecture, which mediates ligand recognition and receptor–receptor interactions^5,7,8^. The extracellular DUF26 domains are also found in plasmodesmata-localized proteins (PDLPs) and cysteine-rich repeat secreted proteins (CRRSPs)^9–11^. Consistent with activation mechanisms established for other RKs, CRKs are proposed to oligomerize and engage extracellular ligands to trigger intracellular signalling. However, to date, there are very few studies demonstrating interactions among different members of the CRK family. For example, CRK28 forms homo- and hetero-oligomers with CRK29, yet the mechanistic consequences of these interactions for specific immune or developmental outputs remain unresolved^12^. Overall, CRK ligands and interaction partners are unknown, limiting functional insight into this family.

Computational biochemical and structural prediction provides a tractable route to infer RLK architecture, potential interaction interfaces, and dimerization modes, thereby guiding hypothesis generation and experimental design^13,14^. Advances in deep-learning structure prediction, exemplified by AlphaFold, now permit high-accuracy modelling of individual proteins and of protein–protein assemblies, offering atomic-level hypotheses for binding interfaces and structural compatibility^15^.

Although predictive models do not replace biochemical and genetic validation, they enable prioritization of experiments and interpretation of sequence–structure–function relationships.

Here, we combine population-genetics and structure-prediction approaches to characterize the Arabidopsis CRK family. Using a panel of 69 natural accessions, we reconstructed the CRK phylogeny and confirmed five well-supported phylogenetic clades. Within each clade, we quantified nucleotide diversity (π) and performed selection neutrality tests, revealing heterogeneous diversity profiles across CRKs and signatures of both purifying and balancing selection acting at different loci. To explore potential functional determinants, we analysed sequence features and AlphaFold-predicted structures of CRK ECDs. Predicted CRK ECD folds are broadly similar to experimentally determined DUF26-containing structures (PDLP5, PDLP8, and ginkbilobin-2) but show distinct glycosylation patterns and disulfide-bond topologies. We further generated pairwise CRK-ECD dimer models (780 combinations); AlphaFold Multimer produced 145 high-confidence dimer predictions, of which ∼78% adopted a conserved dimer conformation. Many predicted dimers form an extended intermolecular β-sheet at the interface, suggesting a putative general interaction mechanism. These results define structural and evolutionary features of CRK ECDs that prioritize candidates and interfaces for biochemical and genetic follow-up to elucidate CRK-mediated signalling. Moreover, this work offers a range of computational approaches to explore biochemical and structural features of the protein at the family level.

## Results

### CRKs nucleotide sequence diversity, dynamics and phylogeny

Nucleotide diversity is a key feature to examine, as it offers insights into the evolutionary processes shaping genes—particularly those involved in vital functions such as development and defence responses^16,17^. Despite the CRK family in *Arabidopsis thaliana* comprising 44 members, in our studies, we focused on 40 CRKs^2^, as the excluded CRKs lacked either an extracellular or a kinase domain or appeared to be pseudogenes (Supplementary Table 1, Supplementary data 1). We assessed the natural genetic variation of CRKs across 69 naturally occurring *Arabidopsis thaliana* ecotypes and obtained 2,760 CRK sequences corresponding to 40 CRK members (Supplementary data 2). Phylogenetic analysis of the evolutionary history of CRKs revealed that they cluster into the five major groups, independent of ecotype or sampling location of origin (Figure 1A). The following groups were defined: (i) group 1 (CRK1, CRK2, CRK3 and CRK42); (ii) group 2 (CRK26, CRK27, CRK28, CRK29 and CRK41); (iii) group 3 (CRK36, CRK37, CRK38, CRK39 and CRK40); group 4 (CRK11, CRK12, CRK13, CRK14, CRK16, CRK17, CRK18, CRK21, CRK22, CRK24, CRK30, CRK31, CRK32, CRK33 and CRK34); group 5 (CRK4, CRK5, CRK6, CRK7, CRK8, CRK10, CRK15, CRK19, CRK20, CRK23, CRK25). As previously reported by Vaattovaara et al. 2019, Group 1 resembles the so-called basal clade. The genes encoding basal clade CRKs were shown to be spread across multiple chromosomes and likely evolved early, with orthologues present in all vascular plants. ^18^ The remaining groups 2-5 constitute the variable clade CRKs, and they are mostly clustered on Chromosome 4 in Arabidopsis and are less conserved, likely having diversified through the whole genome duplication events ^18^. We examined these five CRK phylogenetic groups and analysed the variation in genetic diversity and segregating alleles.

**Figure 1.**
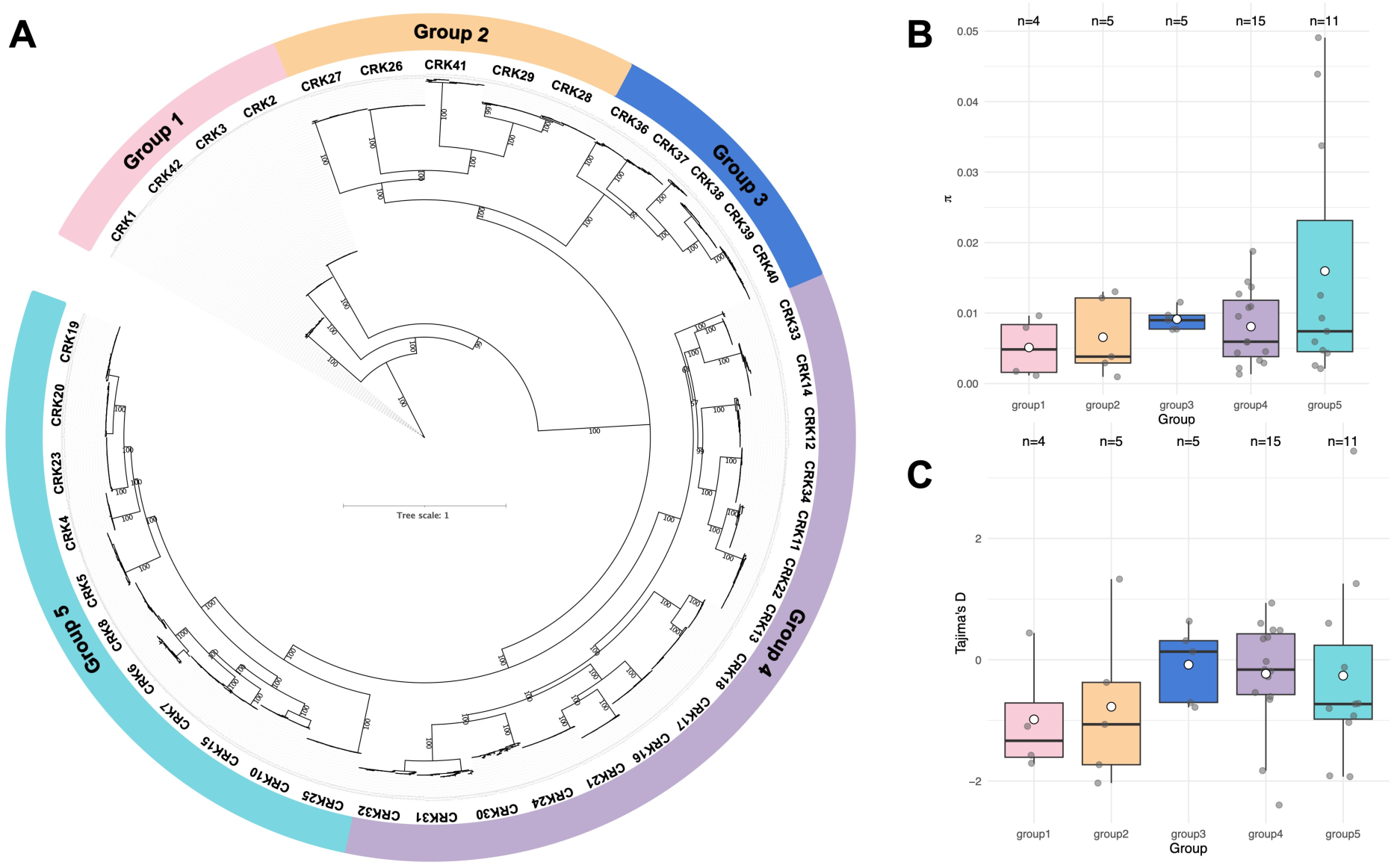
CRK phylogeny and natural genetic diversity. A) Phylogenetic reconstruction based on the CRK coding sequence region across Arabidopsis ecotypes. Best model fit according to BIC score: TPM3u+F+I+R4. Numbers correspond to bootstrap values generated from 1000 ultrafast bootstraps. B) nucleotide diversity (π) and C) Tajima’s D per CRK grouped per phylogenetic group. White dots correspond to the mean, and light grey dots correspond to individual CRK members belonging to that group. Phylogenetic groups represented in the x-axis, n corresponds to the number of CRKs per phylogenetic group.

To understand the evolutionary history of CRKs, we estimated sequence-level measures of genetic diversity — π (nucleotide diversity) and Tajima’s D — for each CRK (Figure 1B and 1C). The index π reflects the amount of nucleotide differences when pairwise comparing all sequences within a phylogenetic group, with a higher π indicating higher genetic diversity. Tajima’s D was used as an indicator of whether a sequence is evolving globally under neutral evolution (primarily stochastic accumulation of mutations) or non-neutral evolution (i.e., natural selection shaping the distribution of mutations). When Tajima’s D is above or below a value of +2 or -2, respectively, that indicates non-neutral evolution at the whole sequence level. We obtained π and Tajima’s D for each CRK and compared them across all five phylogenetic groups (Figure 1B and 1C). Group 5 showed the highest levels of genetic diversity (π=0.0178±0.0190), followed by group 3 (π =0.0091±0.0016), group 4 (π =0.0081±0.0052), group 2 (π =0.0066±0.0056), while group 1 showed the lowest values (π =0.0051±0.0043). These π values indicate that CRKs accumulate genetic variation differently. For Tajima’s D, we found that the mean values across all groups were negative. Tajima’s D ranged from - 0.984 in group 1 to -0.081 in group 3. While negative values of Tajima’s D can reflect an excess of rare alleles, we cannot disentangle neutral or non-neutral evolution at the whole-sequence level, as no group showed values above +2 or below -2. However, when considering individual CRKs, CRK10 had a value of 3.44, while CRK26 and CRK24 had values of -2.03 and -2.40, respectively, indicating non-neutral evolution (Supplementary Figure 1). The values for genetic diversity and Tajima’s D that we found for the entire CDS region of CRKs are in the range of nucleotide-binding leucine-rich repeat (NLR) receptors in rice and A. thaliana^19,20^ .

To further explore the genetic variation in the CDS region, we identified haplotypes within each CRK group (Supplementary Figure 2). A haplotype corresponds to at least two CRKs having an identical amino acid sequence for the entire CDS region. In group 1, from a total of 276 sequences, 200 were distributed in 38 haplotypes (72%). In group 2, out of 339, 267 sequences were placed in 41 haplotypes (72%), in group 3 out of 254, 172 sequences were placed in 46 haplotypes (67%), in group 4 out of 852 sequences, 684 sequences were placed in 128 haplotypes (80%) and in group 5 a total of 504 sequences out of 623 were placed in 102 haplotypes (81%). The most common CDS haplotype among all CRKs was CRK24 from ecotype Abd-0, which was identical in 36 ecotypes. Taken together, the variation in the coding regions of CRKs and the grouping into haplotypes might reflect an interplay between abiotic and biotic factors distributed across different geographical regions.

### CRK domain structure and diversity

To examine the diversity of the CRK family members in more detail, we considered the amino acid composition and domain structure of CRKs (Supplementary Figure 3). The pairwise identity of the amino acid sequence of the full-length protein is higher than that of the ECD alone, indicating that the ECDs are more variable than the rest of the protein (Supplementary Figure 3B, Supplementary Figure 4). This highest diversity in the amino acid composition on the level of ECD can be explained by the general role of the ECD in other RKs families in the perception of diverse signalling molecules and complex formation with other RKs^21^.

Advances in structure-prediction software, such as the development of AlphaFold (AF), enable reliable structural prediction of many proteins, including the individual domains of CRKs. The quality of AF predictions can be measured by the pLDDT (predicted local distance difference test) score. pLDDT scores can be ranked as >90 for very high confidence predictions, 70-90 for high confidence predictions, 70-50 for low confidence predictions, and <50 for very low confidence predictions. However, AF is unable to model full-length single-pass transmembrane proteins (Supplementary Figures 5A, 5B, and 5C) ^22,23^. Thus, we focused solely on the ECD models in our analysis. AF models of CRK-ECDs (excluding the flexible juxtamembrane region) from the selected representatives of each phylogenetic group consistently yielded very high pLDDT scores (>90) and showed a similar fold for all CRK-ECDs, which suggests that AF accurately predicts the CRK-ECD structures (Supplementary Figure 5D). The predicted CRK-ECDs consist of two DUF26 domains. Each DUF26 domain has two α-helixes connected to an antiparallel β-sheet containing five strands (Supplementary Figure 5D). The two DUF26 domains are connected by a flexible loop and through hydrophobic residues in their β-sheets.

To gain further insight into the putative function of the CRK family members, we analysed in detail the predicted structures of the CRK EDCs. We investigated homologs of CRK ECDs to determine whether CRKs could function similarly. DUF26 domains are present in two other families of proteins: Plasmodesmata Localising Proteins (PDLP), and Cysteine-Rich Receptor-Like Secreted Proteins (CRRSPs). The structures of three DUF26-containing proteins, the CRRSP Ginkbilobin-2 (GNK2), PDLP5, and PDLP8, have been experimentally resolved ^18,24^.

PDLPs have an ECD consisting of a tandem of DUF26 domains and a TMD similar to CRKs, but they lack a kinase domain. CRK ECD AlphaFold models closely resemble the PDLP5 and PDLP8 crystal structures ^18^. PDLP-ECDs are the most similar to basal clade CRKs, and previous studies reported PDLPs have likely evolved from basal clade CRKs (Figure 2A) ^18^. PDLP5 aligns to CRK42 with a root-mean-square deviation (RMSD) of 3 Å, indicating that the backbone atoms in the aligned structures are, on average, 3 Å apart. Both PDLPs and basal clade CRKs have 12 conserved cysteine residues forming six disulphide bonds, whereas variable clade CRKs display divergent cysteine conservation. PDLPs have no known ligand. However, they were shown to be involved in regulating callose deposition at the plasmodesmata ^25^.

**Figure 2.**
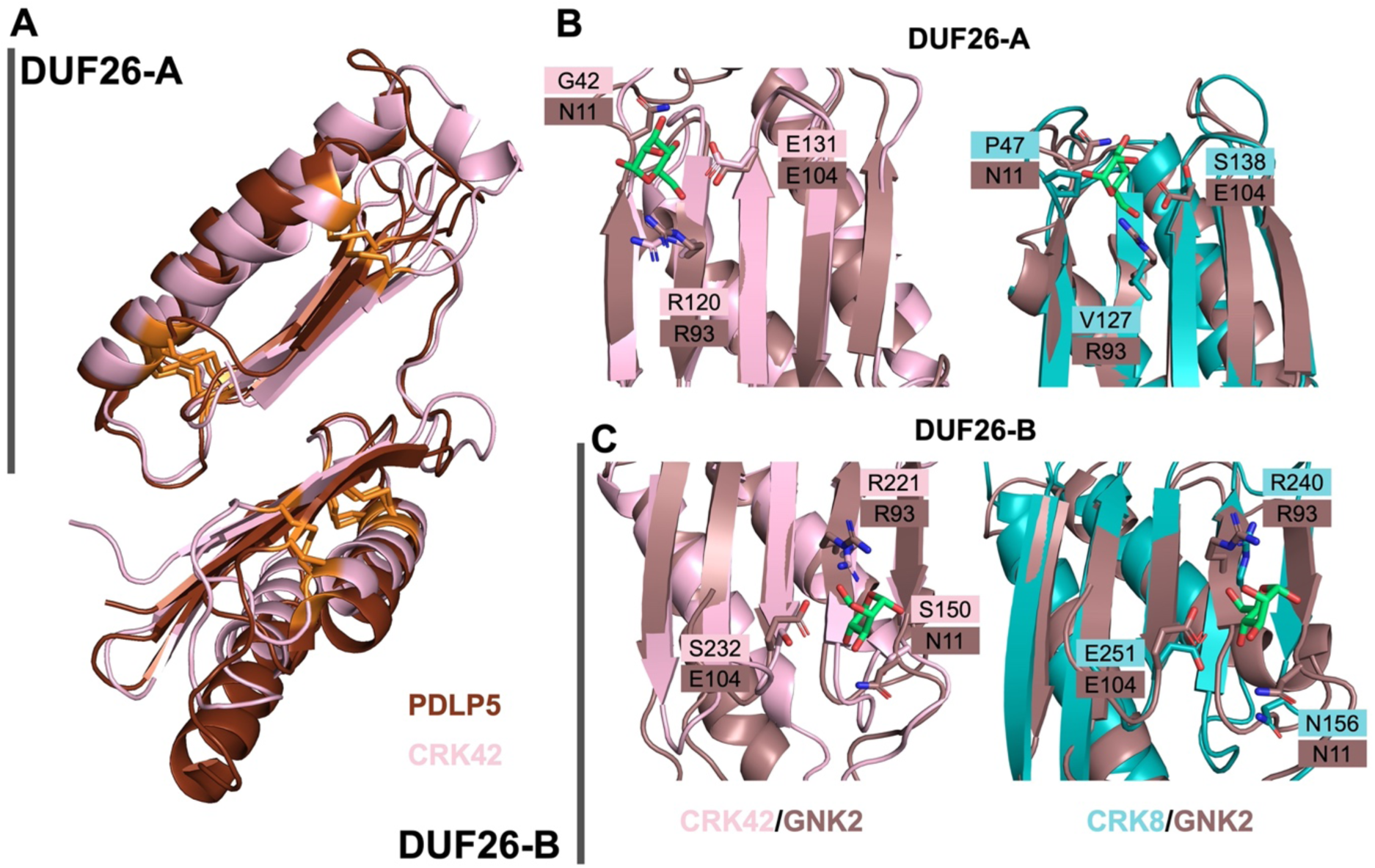
Structural alignments of CRK-ECDs and their homologs. A) Structure alignment of PDLP5 (Brown, PDB: 6GRE^18^) to CRK42-ECD (Pink). Group-1 CRKs are most similar to PDLPs, including 12 conserved cysteines (marked in orange). Structure alignment of GNK2 (brown, PDB: 4XRE^24^) to B) CRK42-ECD (pink) and CRK8-ECD (teal) DUF26-A and C) CRK42-ECD and CRK8-ECD DUF26-B. Mannose is indicated in green. The three residues involved in mannose-binding by GNK2 (N11, R93 and E104) and the respective AA in a similar position in the CRKs are represented as sticks.

GNK2 is a single DUF26 domain-containing protein from *Ginkgo biloba* with anti-fungal properties^24,26,27^. The structure of GNK2 has been resolved by both X-ray crystallography and NMR spectroscopy ^24,27^. Antifungal properties of GNK2 are most likely obtained due to its ability to bind mannose via residues N11, R93 and E104 ^24^. Sequence alignments of GNK2 to CRK ECDs show GNK2 is most similar to the DUF26-A domain of Group-1 (basal clade) CRKs, and 2 of the mannose-binding residues are conserved (Figure 2B, Supplementary Figure 6). However, for variable clade CRKs, no mannose-binding residues are conserved in DUF26-A. Interestingly, if GNK2 is aligned only with the DUF26-B domain of CRKs, it is shown that all three mannose-binding residues, although not all residues in the binding pocket, are conserved in Group-5 CRKs (Figure 2C, Supplementary Figure 6). In addition, other CRK-family members have one or two GNK2 mannose-binding residues conserved. This suggests that certain CRKs may act as glycan-binding proteins.

Although the fold of all CRK-ECDs is similar, CRKs are expected to be involved in distinct signalling pathways, as are other RKs. By analysing the diversity of amino acid sequences combined with the predicted structural data of CRK-ECDs, certain features were identified that may offer valuable insights into protein function. We generated amino acid sequence logo plots of the CRK-ECDs for each phylogenetic group, using the predicted structures of the representative CRKs as a reference, highlighting conserved and variable regions (Supplementary Figure 7). Conserved regions, such as the C-X8-C-X2-C motif, are hallmarks of this family of receptor kinases and are found across all phylogenetic groups. Additionally, each phylogenetic group has a unique set of conserved and variable regions, as shown in Figure 3. Especially those regions can indicate functional diversification.

**Figure 3.**
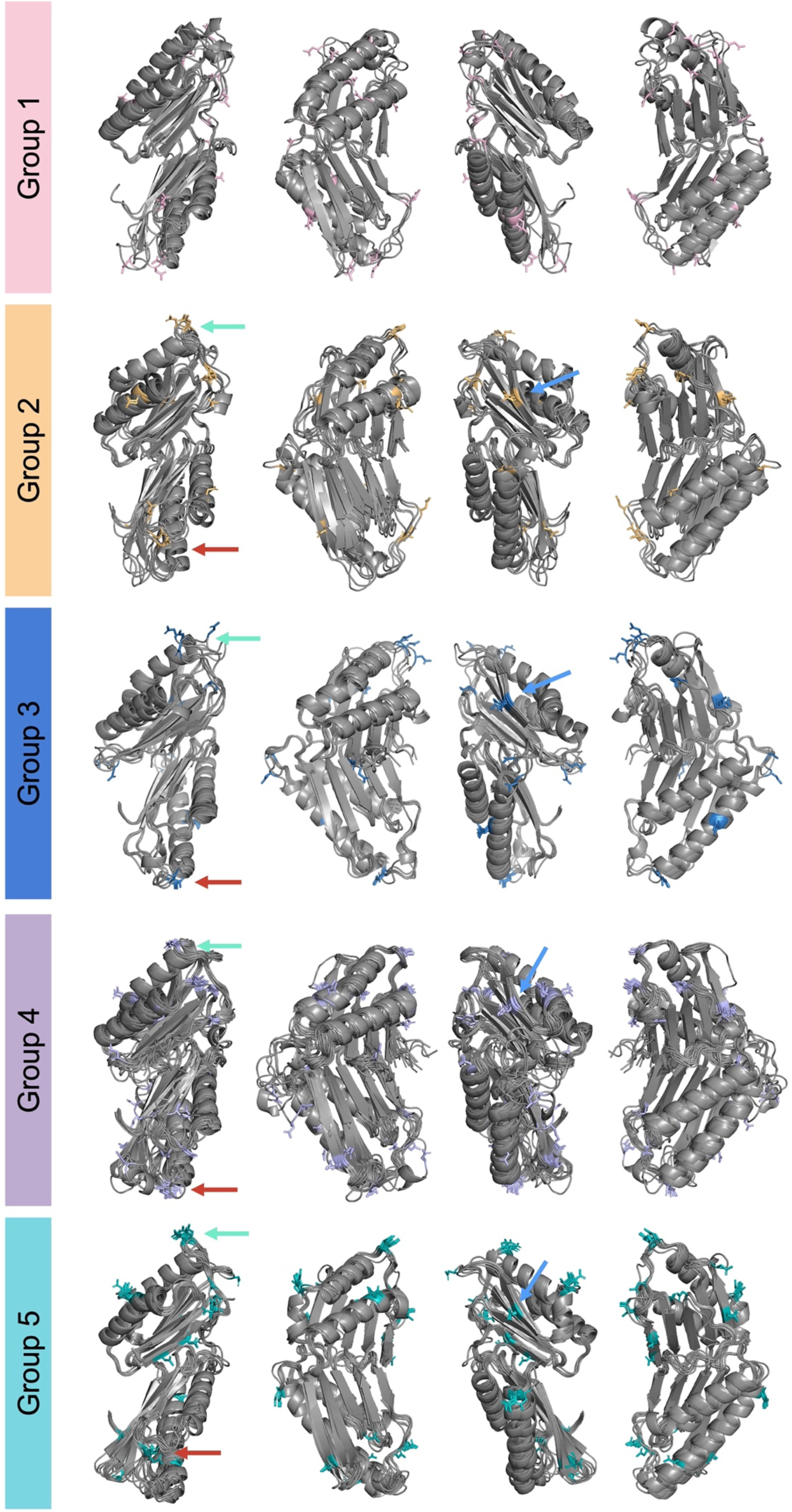
CRK ECDs have multiple glycosylation sites. Predicted N-glycosylation sites of aligned CRK-ECDs per phylogenetic group based on the N-X-S/T motif (where X is not Pro). Asparagine residues that can be glycosylated are coloured by phylogenetic group colour. Three regions were identified (green, red and blue arrows), where CRKs have a conserved glycosylation site.

Finally, we tested for signatures of positive and negative selection in representative CRK sequences from each group. We found signatures of both positive and negative selection (Supplementary Figure 8). In CRK39, we found a total of 34 amino acid sites under selection; in CRK29, 30; in CRK8, 30; in CRK42, 29; and in CRK11, 20. From the identified sites under selection, 47% (16 out of 34) were in the ECD region of CRK39, 50% (15 out of 30) in the ECD region of CRK29, 37% (11 out of 29) were in the ECD region of CRK42, 25% (5 out of 20) in the ECD region of CRK11, and 23% (7 out of 30) in ECD region of CRK8. Together, these results highlight the contribution of different selective forces in shaping the natural genetic diversity of CRKs in A. thaliana. Considering the CRK ECD structures for the analysed representative CRKs, it is apparent that most of the residues under the selection pressure are located in the alpha helices of either DUF26 A or B. Fewer residues under selective pressure are located in beta-sheet regions. Interestingly, there was a clearly higher contribution of the negative selection pressure. Negative selection acts on decreasing genetic variation that reduces fitness, for instance, removing mutations that would impair the sensing of pathogenic organisms or environmental cues. Some of these regions or residues contribute to the functional characteristics of the CRKs’ ECDs, including protein homology, glycosylation sites, cysteine residues, and probability for dimerization. In the following sections, we will explore these characteristics in greater detail.

### Glycosylation of the ECD is likely crucial for correct CRK expression and function

Glycosylation is essential for the folding and stability of proteins but can also be crucial for their function, interaction, and cellular processes localisation^28–30^. de Oliveira et al. 2016 showed that glycosylation of CRK4 and CRK5 was required for BAK1/SERK4-mediated cell death ^31^ and suggested that glycosylation of CRKs is required for the folding of the protein, similar to other RLKs ^30^. The type of sugar group and its attachment vary between glycosylation sites and organisms. Generally, two types of glycosylation are distinguished based on the attachment to the protein: N-glycosylation and O-glycosylation. O-glycosylation sites and the type of glycans attached cannot be predicted using the available predictive tools due to their complexity. However, potential N-glycosylation sites can be predicted for CRK-ECDs (Figure 3). We performed these predictions for all the members of the five phylogenetic groups. When the CRK-ECD models are aligned within phylogenetic groups, three regions stand out in which almost all representative CRKs have predicted N-glycosylation sites (Figure 3). These regions are the relatively flexible loops of DUF26-A and DUF26-B (marked in green and red, respectively). Glycosylation at these positions might be important for protecting the most exposed part of DUF26 domains from, for example, proteolysis or oxidation. The third region (marked in blue) has a predicted glycosylation site on the side of DUF26-A and could potentially be used to shield the exposed edge β-strand, which can be prone to aggregation ^35^. Only the basal group 1 shows no clearly discernible glycosylation pattern among CRKs in this phylogenetic clade.

### The potential functional role of cysteine residues in CRKs

One hallmark of CRK-ECDs is their number of cysteine residues (between 10 and 13), of which most are highly conserved (Figure 2, Supplementary Figure 7). When clustering CRK amino acid sequences by phylogenetic group, a distinct pattern of cysteine conservation emerges. In the basal clade (Group-1), CRKs have 12 conserved cysteines. However, the variable clades (groups 2-5) diverge from this pattern and the number of cysteines. Notably, Group 5 entirely lacks the two cysteine residues at positions 167 and 252, while Groups 2 and 4 lack the cysteine at position 167 and instead have two sequentially adjacent cysteines at positions 252 and 253 (Supplementary Figure 4, Supplementary Figure 7). Lastly, Group-3 CRKs have several cysteine residues that are variably positioned. These variations in the number and positions of cysteine residues between the basal clade and the variable clades could suggest functional divergence.

Based on the protein crystal structure of PDLP5/PDLP8 and AF predictions, most cysteine residues in DUF26 domains are likely involved in disulphide bond formation (PDB: 6GRE, 6GRF)^18^ (Figure 4A). However, this does not apply to non-conserved cysteine residues. Compared to the basal clade, Group-1 and Group-5 CRKs lack a disulphide bridge, Group-3 CRKs often have an additional free cysteine, and CRKs in groups 2 and 4 have two sequentially adjacent cysteines (Figure 4B). AF models suggest that the adjacent cysteines may form vicinal disulphide bonds. Vicinal disulphides are relatively rare and are often found in their reduced form or in disulphide bonds with other cysteines (instead of with each other) ^32^. If the vicinal cysteines in CRKs do form a disulphide bond, as is predicted by AF, then they are likely to have a functional role.

**Figure 4.**
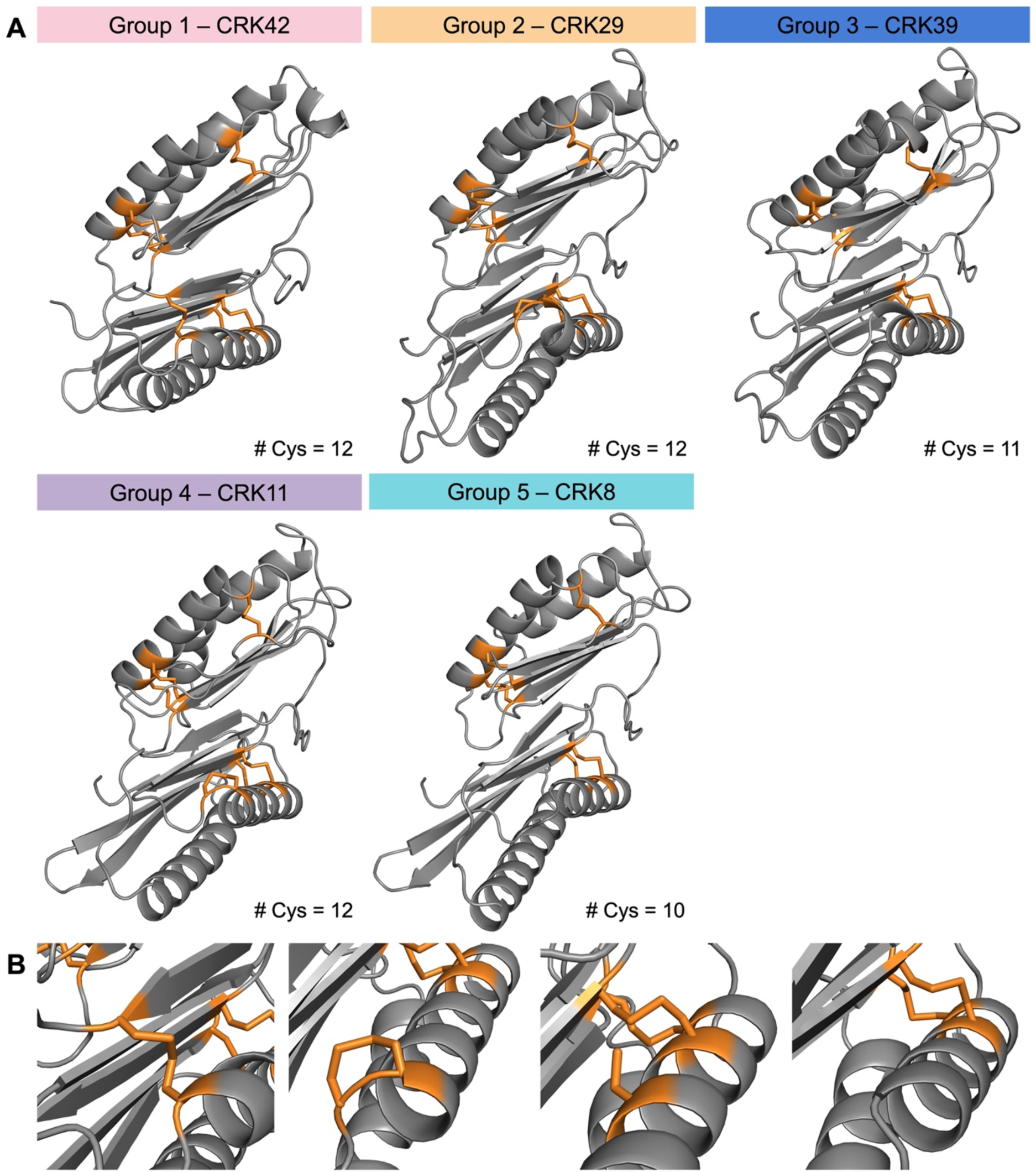
The distribution of the cysteine residues in CRK-ECDs. A) Predicted models of representative CRKs for each phylogenetic group. The cysteines are marked in orange. The representative CRKs were chosen based on the highest average amino acid pairwise percentage identity of each CRK-ECD to the other members of the same phylogenetic group (Supplementary Figure 3). B) The non-conserved cysteines are positioned on the first β-strand in DUF26-B of CRKs. There are four possible options in this position, from left to right: a disulphide bridge connecting secondary structural elements (Group-1), a vicinal disulphide bridge (Group 2 and 4), a single cysteine (Group-3), or no cysteine (Group-5).

### Predicted mode of CRK-ECD dimerization

A commonly reported mechanism for RK signal perception and transduction is dimerization of their extracellular domains. Few studies have reported CRK dimerization. Yadeta et al. showed that CRK28 dimerizes with itself and with CRK29 at the full-length protein level ^33^. To establish a molecular basis for CRK interaction, we predicted CRK-CRK ECD dimer models using AlphaFold2 ^34^. Of the 40 CRKs, we used 38 CRK-ECD sequences as input, excluding CRK23, which has three DUF26 domains in its ECD, and CRK24, which only has one. We predicted pairwise dimer models, generating 741 interaction pairs. For each CRK-ECD pair, five models were generated. The quality of these models was assessed using two metrics: the interface Predicted Template Modelling score and predicted Template Modelling score (ipTM+pTM), and the calculated “interface Predicted Aligned Error” (iPAE) score (Figure 5A and B). The ipTM+pTM score is a metric that indicates the overall quality of the predicted structure, ranging from 0 for low-confidence predictions to 1 for high-confidence predictions ^34^. The PAE is the expected error in the relative positions of residues towards each other, measured as the distance in Angstroms (Å). For the iPAE score, we averaged the PAE scores across domains. For each pair, the best (out of 5) model that met our cut-offs (ipTM+pTM > 0.8 and iPAE < 6 Å) was selected, leaving 174 pairs as high confidence (Figure 7C). We then used PDBePISA to further characterize the interface of all 174 remaining pairs, generating the number of interface residues per model in the pair. Each interface had, on average, 34 +/-5.5 residues per model. A higher number of residues in the interface may indicate a stronger interaction, with more potentially interacting residues. We used the number of interface residues to set a new cut-off both models require 29 (average – 1 standard deviation) or more interface residues, leaving 145 (20%) of the initial CRK-ECD pairs (Figure 5D, Supplementary table 2). Notably, the number of CRK-ECD pairs predicted with high confidence varies between phylogenetic groups (Figure 5E).

**Figure 5.**
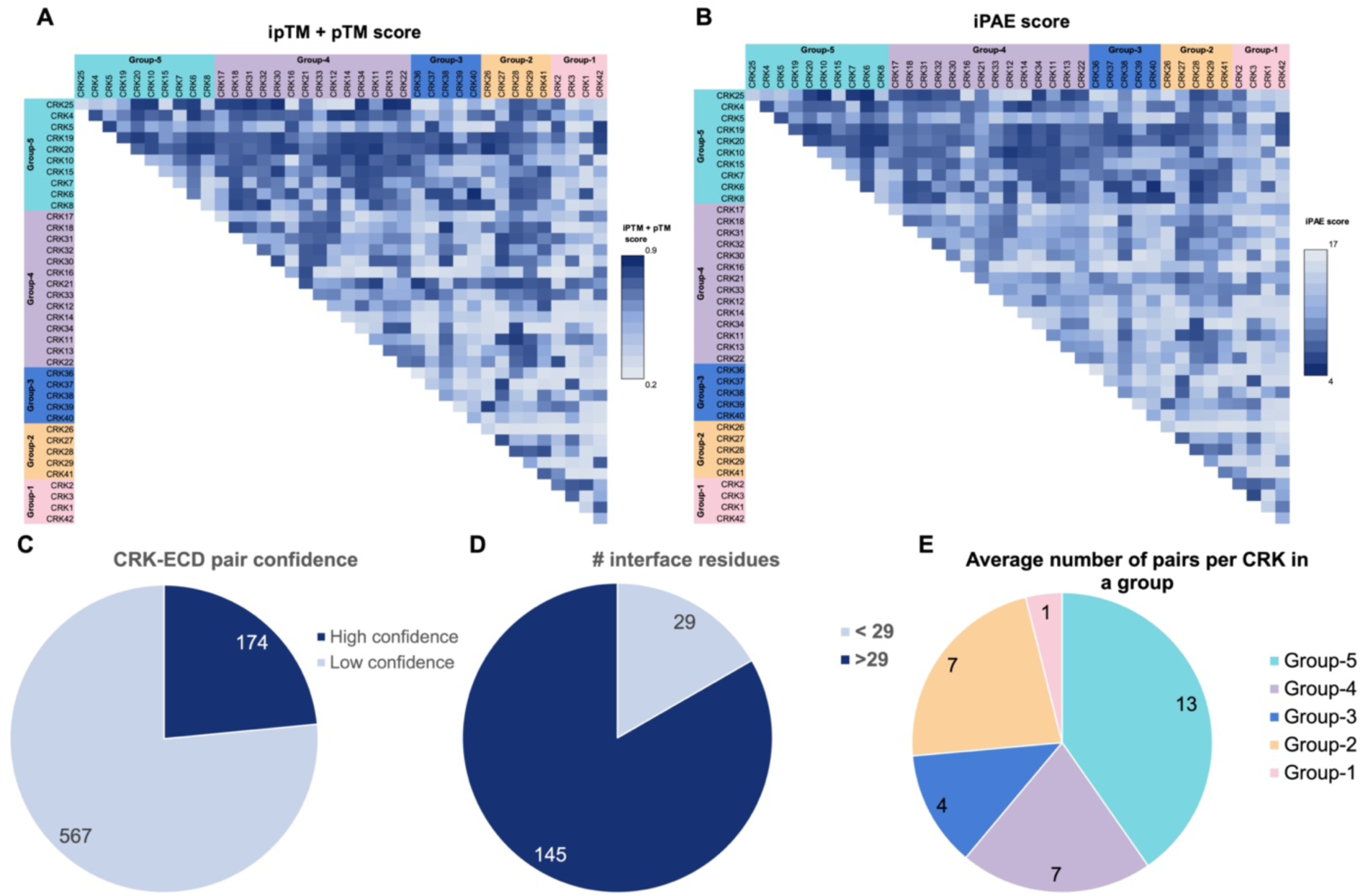
AF predicts the dimerization of CRKs. Heat maps of A) ipTM+pTM and B) iPAE scores for all CRK-CRK predictions. C) The number of CRK-ECD pairs, out of all 741 pairs tested, that pass our ipTM+pTM, iPAE score cut-offs D) A pie chart representing the CRK-ECD pairs out of 174 that have more or less than 29 residues in the interaction interface suggesting high-confidence models. E) Average number of CRK-ECD pairs that pass cut-offs per number of CRKs in a group.

CRK-ECD pairs selected, the ECDs may be oriented differently relative to each other. Three orientation models can be distinguished, which we named: Standard, Flipped, and Flipped and Shifted (Figure 6A). In 78% of high-confidence dimer pairs, the ECDs are in the Standard conformation (Figure 6B). In this conformation, both ECDs are oriented so that the same side of each ECD interfaces with the other. The first β-strand of DUF26-B on each ECD forms the core of the dimerization interface. In the remaining 22% of pairs, the ECDs are in one of two other orientations (Flipped or Flipped and shifted). In 14% of models, one ECD is flipped with respect to the other. In the Flipped orientation, DUF26-A of one ECD faces upward, while in the other, it faces downward, and the first β-strand of DUF26-A faces the first β-strand of DUF26-B of the other monomer. Finally, 8% of the models are in a Flipped and shifted orientation, where one ECD is flipped with respect to the other and shifted vertically, the same β-strand of DUF26-B of each ECD still interacts. If we analyse the distribution of the orientation types per member from each phylogenetic group, Group-1 and Group-3 have few dimer predictions in the Standard orientation, while in Group-2, Group-4, and Group-5 the majority of dimers are in the Standard orientation (Figure 6C).

**Figure 6.**
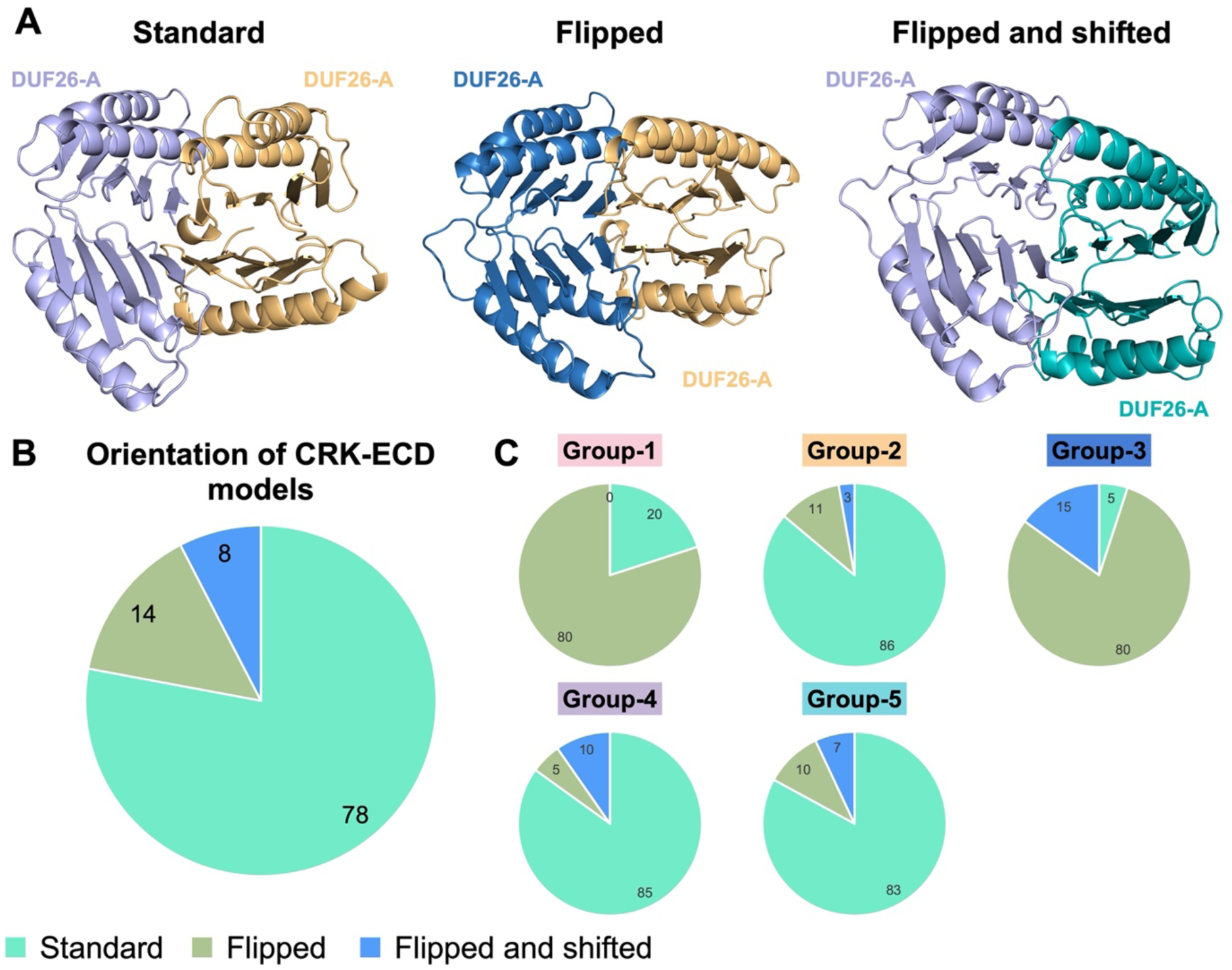
Orientation and distribution of predicted CRK-ECD dimers. A) Three different classifications of orientations were identified, left to right: Standard, Flipped, and Flipped and shifted. The CRK pairs modelled are from left to right CRK31-CRK27, CRK38-CRK27, and CRK11-CRK7. The DUF26-A position is marked, indicating the orientation of the ECD. B) The percentage of each orientation that occurs in our data sets. CRK dimers are in the Standard orientation in 78% of the models. Other orientations occur less often. C) Occurrence of each orientation per phylogenetic group. Groups 2,4 and 5 all have a similar frequency at which the four orientations occur. Groups 1 and 3 have very few models in the Standard orientation.

### The characteristics of the Standard interaction interface between ECDs

In 78% of the high-confidence CRK-ECD interaction pairs, the orientation of the ECDs with respect to each other is in the same Standard orientation, making it a likely candidate for a biologically relevant CRK-CRK dimerization mechanism. To gain further insight into the potential interaction mechanism, we evaluated the properties of the residues and the types of interactions involved in the dimerization interfaces of CRK-ECD pairs in this conformation.

In the Standard orientation, DUF26-B in each ECD form an extended β-sheet together (Figure 7A). In some interaction pairs, the DUF26-A domains of each CRK are also positioned close together, for example, in the CRK27-CRK31 dimer model (Figure 7B). The core of the dimerization interface consists of hydrogen bonds between the first β-strand on DUF26-B of each CRK-ECD partner (Figure 7C). Dimerization through β-sheet extension is common, as the interaction between β-strands from two distinct β-sheets can form similarly to those within sheets ^35^. The residues in this β-strand are not conserved among CRKs, but generally, there are more hydrophobic residues present, which could be an additional stabilising force during dimerization as interaction through hydrophobic patches is a common mechanism (Figure 7D) ^36^. An important factor in protein interaction is the number of contact points, such as hydrogen bonds and salt bridges, between the two molecules. Aside from the main interface through the DUF26-B β-strand, other regions might be involved. The number of contact points varies between CRK pairs and requires further characterisation.

**Figure 7.**
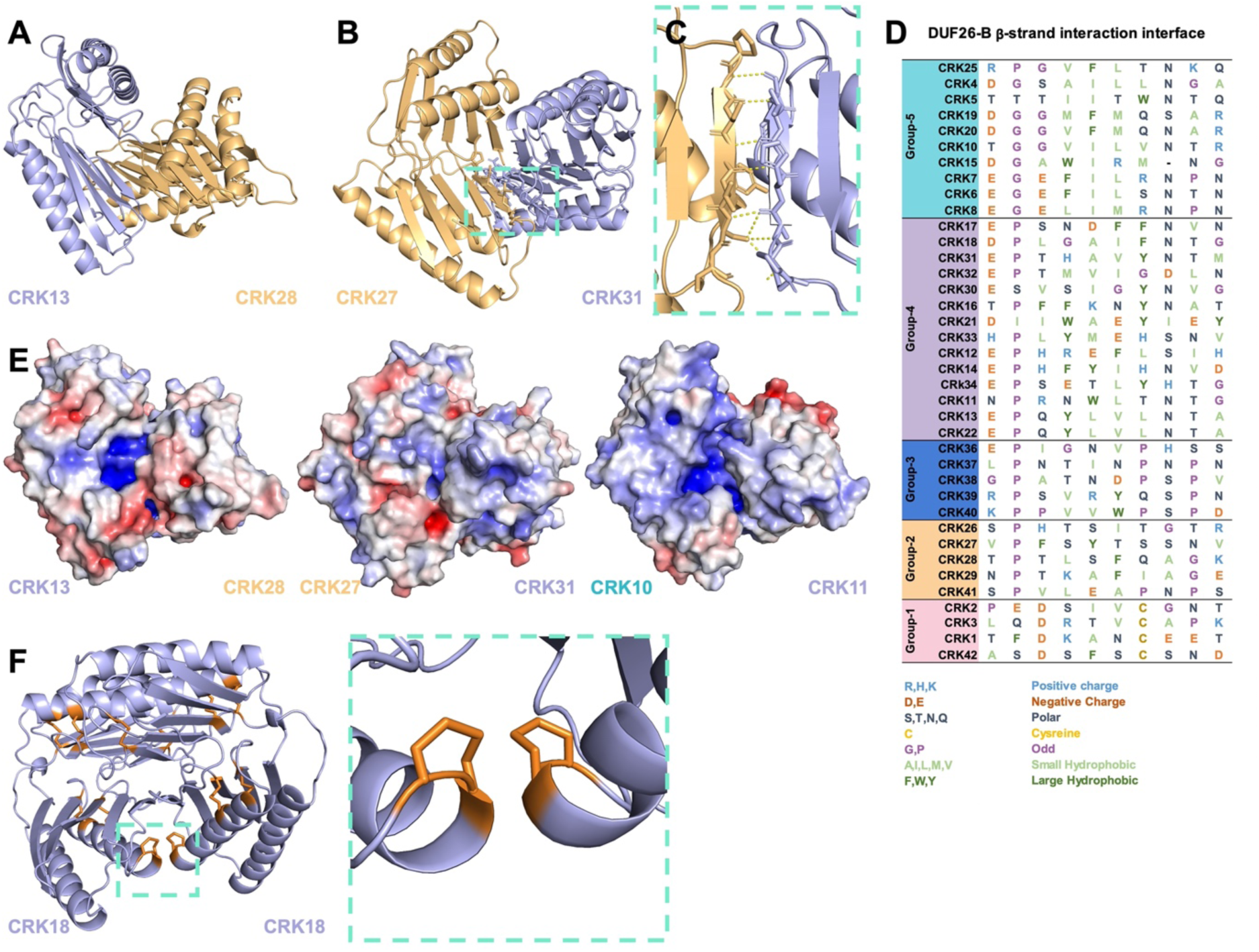
Properties of the various CRK-ECDs interaction interface. A) 72% of CRK-ECD pairs have a predicted dimer interface on the β-strand of DUF26-B of both ECDs. In most Standard orientation models, the DUF26-A domains remain further apart. B) In a small subset, DUF26-A is also predicted to be in close proximity. C) Zoom in on the DUF26-B β-strand interaction shown in (B). The β-strands mainly form hydrogen bonds through the amino acid backbone. D) Amino acid sequence alignment of the interface β-strand. The residues in the interface are not conserved but are more hydrophobic. E) Surface charge predictions of three models of CRK-ECD dimer pairs show surface charge patches surrounding and inside the interface. Surface charge is represented by a colour gradient from red (negative electrostatic potential) to blue (positive electrostatic potential). APBS electrostatics was used in PyMol to generate the maps 26. F) CRK18-CRK18 pair showcasing the proximity of the vicinal disulphides to each other in the dimer. Vicinal disulphides are marked in orange.

Martin-Ramirez et al. 2025 reported that CRK-ECD’s have diverse surface charges, which could contribute to the protein interaction^3^. We selected three interaction pairs, CRK13-CRK28, CRK27-CRK31, and CRK10-CRK11, and modelled the surface charges of the dimers (Figure 7E). In these examples, charged patches also surround the dimerization interface. In addition, in two of the interaction pairs, CRK13-CRK28 and CRK10-CRK11, highly charged patches are present in the interface between the ECDs, which could have a role in the dimerization interaction or ligand binding ^37^.

In the Standard orientation, the vicinal disulphides (present in CRKs from Group- 2 and 4) of each ECD can come in close proximity to each other, for example in the CRK18-CRK18 homodimer (Figure 7F). While the function of the vicinal disulphides in CRKs remains unknown, it is interesting that these potentially reactive residues are brought together by dimerization of CRKs. Taken together the importance of β-strand interactions, surface charges, and vicinal disulphides for dimerization in CRK-ECDs in the Standard orientation provide interesting points for further investigation.

### CRK-ECDs form a highly interconnected predictive interaction network

Interacting receptors are components of complex signalling networks that initially perceive signals and ultimately trigger signal transduction, leading to appropriate cellular responses. To understand the dynamic range of intra-family interactions between CRK-ECDs, network studies were used (Figure 8A) ^38^. In the network, each circle, known as a node, represents a CRK-ECD. The nodes are coloured based on the phylogenetic group to which the CRK belongs. The lines, also known as edges, connect interacting CRK-ECDs and represent predictive physical interactions in this model. The size of the node is relative to the PageRank score, which serves as a centrality measure within the network, where larger node sizes indicate more interactions. Groups of CRKs with common interactors were clustered together into 3 communities (numbered I-III, highlighted by the yellow background) based on the WalkTrap algorithm. Lastly, homodimerization is marked by a grey circle surrounding the node. Homodimerization is predicted for 7 CRKs (CRK27, CRK28, CRK18, CRK4, CRK5, CRK19, CRK20) (20% of the 38 CRKs tested).

**Figure 8.**
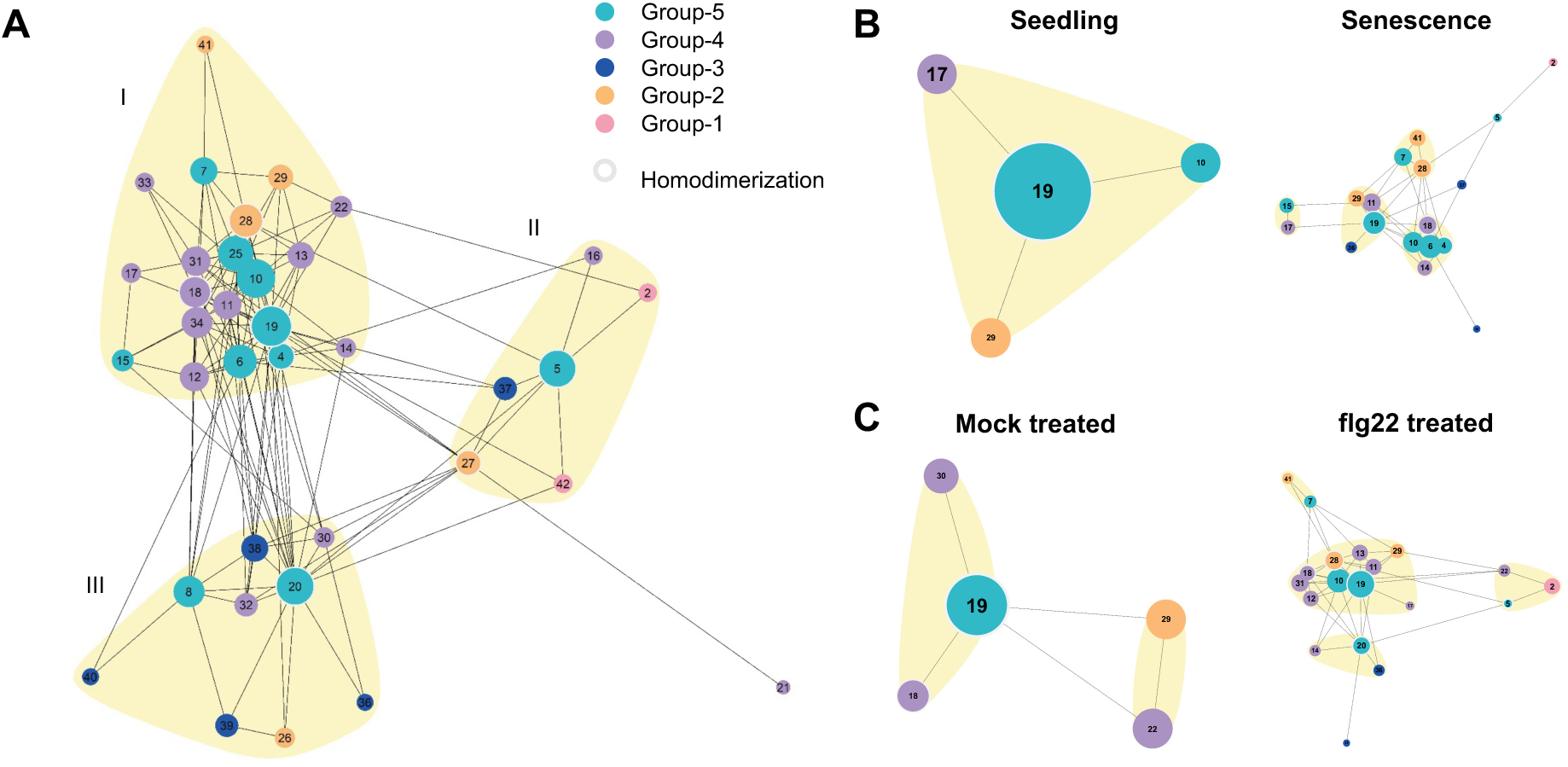
Interaction network of CRK-ECD AF predicted dimers. A) The 145 AF CRK-ECD dimer predictions that passed our cut-offs were used to generate an interaction network. Each number represents a CRK-ECD, the coloured circles or nodes correspond to the phylogenetic group to which the CRK belongs. The node size corresponds to the number of interactions a CRK has. A light grey outline of the node indicates homodimerization. Lines connecting nodes indicate heterodimers. Communities of interconnected CRKs, as identified with the WalkTrap algorithm ^38^, are marked by a yellow background and are numbered I, II and III. B) interaction networks refined using expression data of CRKs in seedlings, senescent plants, and mock-treated or treated with 1 µM flg22.

In general, we observe a high number of connections between CRKs, leading to a dense network (Figure 8A). The Group-1 CRKs form a few interactions that contribute to only one community, which could be a consequence of the different orientations of the ECDs in the dimer preventing interaction with variable clade CRKs. Similarly, the majority of models involving Group-3 CRKs display different dimer orientations and are involved only the smaller communities II and III. The core of the network primarily comprises CRKs from phylogenetic groups 2, 4, and 5. Group-5 CRKs are predominantly represented within the network, consistent with previous observations that Group-5 CRKs have the highest number of interactions (Figure 7E). Interestingly, Group-5 members CRK19 and CRK20, with a high sequence similarity (80% pairwise identity of their ECD), and a high number of interactions, are in different communities in the network. These findings suggest CRK phylogenetic groups may play distinct roles within a larger connected network.

For CRKs to interact, they must be present within the same tissue type at an identical developmental stage. The networks were further refined utilising CRK expression data across various developmental stages, tissue types, and stress treatments sourced from public databases (refs). A higher expression of CRKs is observed in senescent tissue relative to seedlings, as reflected in the interaction networks (Figure 8B). Similarly, Arabidopsis seedlings exposed to 1 µM flg22 exhibit increased CRK expression, leading to a denser network (Figure 10C). In both instances, heightened CRK expression is associated with an increase in ROS production either due to developmental processes, such as senescence, or defence responses triggered by flg22.

The majority of the CRKs found in the filtered networks are found in community I of the global network. It is very clear that seedling and mock-treated seedling networks have a low number of nodes (4-5), which is linked to a low expression of the CRKs at the seedling stage. Seedling and mock-treated seedling networks do not match completely, as they overlap only with CRK19 and CRK29. This could be explained by either biological and experimental variations, as these are two independent data sets or the effects of the mock treatment. While senescence and flg22-treated networks are similar in the number of CRKs (18 and 19 CRKs, respectively) and in the composition of the nodes, the distribution of individual nodes within smaller communities differs. This may indicate that a certain level of specificity is provided by the CRKs that are expressed and interact in response to different stimuli.

## Discussion

The computational predictions at the nucleotide and protein family levels greatly enhance our ability to infer function, understand biological processes, and efficiently guide experimental research. To gain evolutionary, biochemical, and structural understanding, we performed family-wide computational studies, which ultimately can support the experimental direction taken to unravel the biological roles of different CRK members. We surveyed the natural sequence variation, structural features, and predicted interaction landscape of the CRK family.

### Evolutionary dynamics and sequence diversity

The phylogeny identified five well-supported groups that mainly correspond to previous basal-versus-variable classifications but expands on this with group-specific diversity metrics^3,11^ (Figure 1A). Group 5 displayed the highest mean nucleotide diversity (π), whereas Group 1 (basal clade) was the least variable, consistent with an early origin and broader conservation across vascular plants^11^ (Figure 1B). The generally negative Tajima’s D values across groups indicate an excess of rare alleles, but only individual genes (CRK10, CRK24, CRK26) showed Tajima’s D values consistent with deviation from neutrality, suggesting locus-specific departures due to recent selective sweeps, balancing selection or demographic changes (Figure 1C, Supplementary Figure 1). Interestingly, CRK10, which is known to play a role in the activation of defence responses in Arabidopsis, displays significantly higher Tajima’s D than other family members, consistent with balancing selection^39,40^. In this scenario, the alleles are maintained in the population, which is consistent with pathogen virulence genes evolving to evade host immune recognition. Some studies on Arabidopsis plants found low Tajima’s D values for receptor-like proteins (RLPs), similar to those for CRK24 and CRK26, suggesting purifying selection on genes involved in conserved developmental processes and the detection of conserved pathogen-associated molecular patterns (PAMPs)^41^. Haplotype distributions further confirmed that coding variation is unevenly distributed among groups, with some CDS haplotypes widespread across ecotypes (e.g., CRK24-Abd-0) and others more restricted (Supplementary Figure 2). This may be connected to geographically heterogeneous selective pressures or to genetic drift acting on locally adaptive variants^42^. Ngou et al. 2024 also found high levels of nucleotide diversity in genes encoding Pattern-Recognition Receptors (PPRs), indicating strong selective pressures and ongoing diversification, which are vital for adapting to various pathogen populations ^43,44^.

### ECD-centred structural diversification

Amino acid divergence is concentrated in ECDs, whereas the kinase domain is highly conserved, implying the intracellular signalling machinery is maintained while ligand recognition or interaction specificity has diversified^45^ (Supplementary Figures 3, 4, and 5). ECDs of many RLKs bind ligands and interaction partners, making them important determinants of RK function^4,46^. To gain insight into the potential modes of interaction and functions of CRKs, we computationally investigated the properties of their ECDs. We could accurately predict the CRK-ECDs with AlphaFold, yielding models with structures displaying high similarity to the crystal structures of other DUF26-containing proteins, PDLP5, PDLP8 and GNK2 (Figure 2). Despite the high degree of structural similarity, CRK-ECDs are characterized by multiple biophysical and biochemical features, including sequence diversity, surface charge, glycosylation sites, and number and distribution of cysteine residues, that differentiate them from PDLPs and GNK2^3,11,47^. Importantly, many positively and negatively selected sites map to ECDs (23–50% of detected sites in representatives) (Supplementary Figure 8). The predominance of sites in which mutations are underflavored underscores the functional constraints on maintaining key structural elements, such as ligand-binding or interaction partner-binding interfaces. Similarly, in CLAVATA 1 (CLV1), nearly all residues were under negative selection. This was expected, as *clv1* mutants exhibit severe cell proliferation phenotypes across various species, supporting the idea that CLV1 is a conserved developmental gene subject to strong purifying selection^48^. However, CRKs also display localised positive selection, suggesting adaptive tuning of recognition surfaces, likely in response to diverse biotic and abiotic cues, and possibly ROS perception. The work of Kileeg and Mott (2025) identified strong positive selection across many RLK gene subfamilies and a bias of positive selection in the extracellular domains of receptors^42,48,49^. RLKs involved in plant defence responses have signals of positive selection, particularly in regions responsible for pathogen recognition, indicating that these parts of the genes are evolving rapidly to enhance the plant’s immune capacity. This suggests escape from adaptive conflict within the extracellular domain may have played a significant role in the evolution and adaptation of the RLKs. Moreover, positive selection was more common in the LRR domain than in other RLK domains^48,49^. This diversity in the signatures of selection suggests that CRKs may have distinct mechanisms for signal perception, complex formation, and modulation of different signalling pathways, which are essential for plant growth and defence.

### Functional implications of glycosylation and cysteine architecture

Distinct biochemical features can significantly influence the precise regulation of protein function. Therefore, we elected to conduct computational evaluations of some of these features at the family level. In plants, glycosylation is crucial for RLKs’ function, particularly for proper protein folding, trafficking, and stability^50^. It also directly regulates their activity, enabling responses to external stimuli by affecting ligand binding affinity and signal transduction. Key demonstrated roles include maintaining cell-surface receptor function in pattern-triggered immunity (PTI), like for the proper function of EF-TU RECEPTOR (EFR) or developmentally guiding fertilization through pollen tube guidance mediated by FERONIA (FER) and its family members^50–53^. In the CRK family, conserved N-glycosylation motifs cluster in exposed flexible loops of DUF26-A and -B and on an exposed DUF26-A flank. Likely, these glycan moieties would plausibly stabilize ECD folding, protect labile loops from proteolysis/oxidation, and modulate interactions—consistent with prior experimental requirements for glycosylation in CRK function (Figure 3).

Cysteine patterns differentiate basal and variable clades: basal CRKs preserve a 12-cysteine disulfide architecture, while variable clades show gains or losses, including vicinal cysteines (Figure 4). These differences could alter local stability, create or remove redox-sensitive switches, or shape ligand-binding pockets. One hypothesis in the field is that CRK could sense ROS through its cysteine residues ^54,55^. Vicinal disulphides have the potential to be redox active. In mammalian proteins such as von Willebrand factor and BMP-1, vicinal disulphides play key roles in protein function ^56,57^. In these cases, the reduced state of the vicinal cysteines results in a more flexible, less stable protein, a feature crucial for regulating protein-protein interactions. However, Richardson et al. 2017 analysed existing vicinal disulphide-containing proteins in the PDB and showed that redox-active vicinal disulphides are not common^32^. Instead, vicinal disulphides are often involved in ligand binding by providing a rigid hydrophobic contact point for sugars or multiring ligands ^32^. The non-conserved cysteine residues in CRKs may play a functional role. However, if this function involves redox switching, is part of a binding pocket, or has a different role, it remains to be determined experimentally.

### Predicted mechanism of CRK-ECD interaction

Dimerization mechanisms of RLKs can give important insights into to the involvement of the specific CRKs in the regulation of signalling pathways. To gain insight into the possible CRK-ECD dimerization mechanism, we studied the predictive dimerization specificity, dynamics and the orientation of all predicted high-confidence dimers (Figures 5 and 6). In 78% of high-confidence dimers, the CRK-ECDs are in the same orientation. The core of the identified interaction interface is an extended β-sheet formed by the interaction of the β-strands of DUF26-B domains. A similar extended β-sheet is also present in the Flipped and Flipped-and-shifted orientation types. Vaattovaara et al. 2019 proposed a similar dimerization mechanism based on the PDLP5 and PDLP8^18^ crystal structure packing^18^. The extended β-sheet forms an attractive model for a conserved DUF26 domain dimerization mechanism where the properties of the interface may determine specificity.

In general, β-sheets form through β-strand backbone hydrogen bonding. The edges of the β-sheet, also called β-edge strands, could interact further with other β-sheets if left exposed ^35^. Proteins avoid unwanted interaction of β-edge strands through various mechanisms, including strands twisted by proline or glycine, charged residues, short edge strands or by covering edge strands with loops ^58^. In order for β-strands to allow for dimerization, they require 5 or 6 exposed H-bonding residues that are regularly spaced ^58^. CRK-ECDs have four edge strands. The first β-edge strand of DUF26-A is short and poorly predicted. The second edge strand is often partially covered by an α-helix. In addition, DUF26-A has conserved glycosylation sites near its β-edge strands. Glycosylation could further prevent the strands from interacting. The second DUF26-B β-strand is partially covered by a loop. The first β-edge strand of DUF26-B is the most exposed β-strand of CRK-ECDs and is part of the interface in AF predictions.

In addition to the β-strand interaction, the predicted dimers have notable charged regions near the interface (Figure 7). One common mechanism for protein dimerization is through a surface characterised by a hydrophobic patch surrounded by charged residues ^36,59^.

### Dynamics of CRK-ECD interactions

Network analyses of the predicted high-confidence interactions among all CRK-ECDs reveal that the interacting proteins create a dense, highly interconnected network (Figure 8). The communities identified are particularly rich in members of groups 2, 4, and 5. Group-5 CRKs are especially important to the overall connectivity and structure of the networks, as indicated by their high average PageRank score. Among the predicted interacting CRK ECDs, we found that they were previously shown to occur at the level of the full-length receptors homodimerization of CRK28 but not heterodimerization between CRK28 and CRK29^12^. The community I has the highest number of interacting CRK ECDs. Notably, many of them, such as CRK6, CRK7, CRK28, CK29, CRK10, and CRK14, have previously been reported to be involved in defence and abiotic stress responses ^12,60–62^. This is very interesting and may suggest that stress-related CRKs cluster together, possibly forming receptor complexes responsible for stress response signalling.

The formation of these networks depends on CRKs being present in the same tissue and at the same developmental stage, as well as under specific stress conditions. Increased CRK expression during senescence or shortly after stress treatments such as flg22 exposure results in denser interaction networks. The networks also reveal that CRKs are more active and interconnected in certain tissues and under specific stimuli, suggesting functional specialization. Notably, seedling networks with low CRK expression show fewer interactions, while stress or developmental cues lead to complex, expansive networks. Differences in community distribution within similar networks suggest that CRKs may have specific roles depending on the stimulus or stage. Overall, the data emphasize the importance of both phylogenetic grouping and expression patterns in understanding CRK functions within plant signalling networks.

### CRKs as ROS or glycan receptors

Potentially two types of signalling molecules to be sensed by CRKs: ROS and glycans. CRKs have been proposed as ROS receptors because of the high number of cysteines in their ECDs, which could undergo oxidative modification. ROS is produced in the apoplast during stress responses. While it is likely ROS is also perceived in the apoplast as a signalling molecule, the RKs involved in ROS detection remain largely unknown. To date, only the leucine-rich repeat receptor kinase Hydrogen Peroxide-Induced Ca^2+^ increases 1 (HPCA1) has been shown to be modulated by ROS in the apoplast ^63^. CRKs have been linked to many defence responses that produce ROS ^64–66^. It is also known that, transcriptionally, CRK members are upregulated during developmental processes associated with elevated ROS levels or during various abiotic and biotic stresses^3^. However, a direct link between CRKs and ROS has never been demonstrated. Vaattovaara et al. showed that in PDLP5 and PDLP8, all cysteines are involved in disulphide bonds, suggesting a role in the structural stability of DUF26 domains ^18^. However, this assessment does not include the non-conserved cysteines of CRKs. Cysteine residues involved in disulphide bridges are often required for protein stability and as a result conserved ^67,68^, suggesting the non-conserved cysteines in variable clade CRKs may have a functional role rather than a structural one. In addition, the non-conserved cysteines in phylogenetic Groups 2 and 4 are predicted to form vicinal disulphides, which are placed in close proximity in the dimer models. Willems et al. showed AF models of plant proteins correctly predict disulphide bonds ^68^ ^68^. Vicinal disulphides are rare and can adopt unfavourable conformations, making them more likely to be essential for function ^32^. The vicinal disulphides may be susceptible to redox modulation and potential candidates for ROS sensing ^69^.

CRKs may function as glycan receptors due to their homology with GNK2 ^24,26,27^. Glycans can originate from the plant or microbial cell wall and are important signalling molecules in both stress and developmental signalling processes ^70,71^. GNK2 plays a role in fungal resistance through binding of mannose from the fungal cell wall^24^. Other DUF26-containing proteins from maize, AFP1 and AFP2, have also been implicated in glycan binding ^72^ . Additionally, a CRK from wheat, TaCRK3, also showed anti-fungal properties ^73^. Sequence alignments of GNK2 to CRK DUF26 domains show that the mannose-binding motif is conserved in some Arabidopsis CRKs, making them interesting candidates for glycan binding. Recently, the work of Pierdzig et al. 2025 demonstrated that CRK7 functions as a receptor for bacterial-derived wall teichoic acids (WTAs) in Arabidopsis. When CRK7 encounters WTA, it forms homo-oligomers at the plasma membrane —a common mechanism for carbohydrate receptors —activating defence responses^74^.

### The limitations of AlphaFold predictions

AlphaFold offers a model for CRK-CRK ECD dimerization, but several limitations should be considered. AF is trained on experimentally resolved protein structures, and models could be biased toward it. This raises concerns about possible "memorization" from existing PDLP crystal structures.

A clear limitation of AF is its inability to model full-length CRKs, thus it cannot assess if the TMD and KD contribute to the dimerization. Moreover, when modelling interactions of only the CRK-ECDs using AF, we cannot account for the effect of CRKs being anchored in the plasma membrane, limiting its mobility. However, the flexible extracellular juxtamembrane region, which has an average length of ∼35AA in CRKs, could allow CRK-ECDs a large range of motion, allowing them to be in different orientations that fit the predicted dimer configurations. In addition, the “standard” orientation which is predicted in 78% of the high confidence models, would only require each ECD to “bend” to the side with the juxtamemrane region extending from the last β-strand down to the membrane. Interface predictions, while stringent (ipTM+pTM, iPAE, interface residue thresholds), require biochemical and in planta validation: co-immunoprecipitation, Förster resonance energy transfer (FRET), crosslinking mass spectrometry, or cryo-EM/ crystallography of ECD complexes.

In addition, the effects of the full-length protein on the CRK interaction and other factors, such as post-translational modifications (glycosylation), pH, other interacting proteins, and ligands may influence CRK dimerization ^21,75,76^. The recently released AF3 is a step closer to including these conditions, allowing for the prediction of glycosylation and small lipids ^77^. However, it cannot overcome the lack of experimental data needed as input, for example, interaction partners or glycosylation sites of CRKs. While AF can provide valuable insights, it is not a replacement for experimental work ^78^.

### Future perspectives

Our integrated computational studies position CRK ECDs as the principal component of diversification within the family, balancing strong purifying constraints on structural elements with localized adaptive changes in extracellular recognition surfaces. AlphaFold predictions give a likely mechanism for CRK-ECD dimerization, which can be used as a hypothesis to help guide future experiments to elucidate CRK interactions. Further developments in the structure prediction field, such as the recent release of AF3, can provide further insights by including post-translational modifications and ligands in the predictions ^77^. The predicted β-strand–mediated dimerization provides a mechanistic hypothesis for CRK complex assembly, with surface charge, glycosylation and cysteine topology modulating interactions and responsiveness to environmental cues such as elicitor-triggered ROS or cell-wall derived glycans. Moving forward, focused experimental tests are essential: (i) validate key predicted homo- and heterodimers under native expression and membrane conditions; (ii) map glycosylation and redox states and assess effects on folding, localization and complex formation; (iii) probe ligand-binding abilities of DUF26 variants (including mannose-like glycans) suggested by GNK2 homology; and (iv) relate allelic variation to phenotypic outcomes across ecotypes (disease resistance, ROS responses, developmental regulation). Such work will clarify how CRKs incorporate evolutionary variation into plant signalling networks and adaptive responses.

## Methods

### Identification of CRK sequences across naturally occurring Arabidopsis thaliana ecotypes

To identify the nucleotide sequences of CRKs across naturally occurring ecotypes of A. thaliana, we leveraged from the previously published chromosome-level pan-genome of 69 accessions^79^. We used the nucleotide sequences of all available CRKs in the TAIR database to search against the 69 whole genomes (Supplementary data 1). In brief, using makeblastdb (version 2.9.0)^80^. we created local blast databases for each A. thaliana genome. Next, using the nucleotide sequences of each CRK as **a** query (supplementary 2), we performed a local blast search against each genome database. We extracted information on the start and end positions of the best blast hit using a custom R-script. With the identified region in blast we extracted the nucleotide sequences from the target genome with samtools faidx (version 1.10)^81^. Finally, all sequences were concatenated into a single multi-sequence fasta file.

### Nucleotide sequence alignment and phylogenetic tree

The coding region of the extracted CRK nucleotide sequences were aligned with mafft (version 7.525) ^82^ using the "genafpair" option with a maximum of 1,000 iterations. From the resulting alignment, we used trimAl (version 1.5)^83^ to exclude positions with more than 30% of gaps. The final alignment was used to reconstruct the phylogenetic tree with IQ-TREE2 (version 2.4.0),^84^ setting the number of bootstrap replicates to 1000 and automatic evolutionary model search. The final concatenated tree was visualized in iTOL^85^.

### Natural selection, summary statistics and ECD region diversity

For each CRK across haplotypes we removed intronic regions, keeping only the amino acid coding regions (CDS). We generated haplotype networks using the TCS method implemented in PopART package ^86,87^. Additionally, using PopART package we determined the frequencies of haplotypes within each CRK. Nucleotide diversity **(l’l)**(π) and Tajima’s D were computed using the R-packages PopGenome (version 2.7.7) and pegas (version 1.3). We used the HyPhy package^88^ to search for signatures of natural selection considering three underlying models of evolution: Mixed effect model of evolution (MEME)^89^, fast unconstrained Bayesian approximation ^90^ and single-likelihood ancestor counting ^91^. AliView (version 1.26)^92^ was used to translate the nucleotide coding region of our CRK clade-groups into amino acids. After retaining only the extracellular domain, we generated logo plots using the Python module Logomaker (version 0.8.6)^93^.

### Sequence alignments

Amino acid sequences of Arabidopsis CRKs were retrieved from TAIR. The pairwise identity matrix and multiple sequence alignments were generated using Clustal-O (version 1.2.4)^94,95^ and Muscle (version 3.8)^96^ respectively, using the EMBL-EBI job dispatcher^97^.

### AlphaFold modelling

Full-length CRK models were extracted from AF-database ^98,99^. All other AlphaFold predictions were performed using a local installation of AF v2.2.2. Parameters were set to default; five random seeds were generated with one model per seed. For monomer ECD models, the amino acid sequence from the first residue to the predicted start of the transmembrane domain was used as input. The highest scoring models were used for the figures, and N- and C-terminal flexible regions with a low pLDDT confidence score were removed.

To model the CRK-ECD dimers we cropped the input sequence to contain only the structured DUF26 domains, as low confidence flexible N- and C-terminal regions could interfere with the predictions. Dimer models were predicted for 780 pairs, generating five models each. The ipTM+pTM scores were generated by AF. PAE scores were generated by AlphaPickle, and the interface specific scores (iPAE) were extracted^100^. Per CRK-ECD pair the highest scoring model that had an ipTM+pTM score and an iPAE score above 0.8 and an iPAE score smaller than 6 Å was selected as a high-confidence prediction. Next, for each model passing ipTM+pTM and iPAE scores the number of interface residues was determined using PDBe PISA^101^, models with 29 (average – 1 standard deviation) or more residues in each CRK-ECD interface were selected. The 145 CRK-ECD pairs that were modelled with high confidence were further inspected manually.

### Network construction and analysis

The network was constructed using the igraph package (http://igraph.org/r/) in the R programming environment (https://www.r-project.org/). Clusters of interacting proteins in the network were identified using the WalkTrap algorithm as implemented in igraph using a random walk length of 8 (reference DOI: 10.1038/srep05739)^102^. The PageRank algorithm implementation using the PRPACK library within the igraph package was used as a centrality measure within the network, with the node size set to be relative to the PageRank score.

## Authorship contribution

JFS designed the experiments. SM and DdSP ran the AF predictions. DdSP and JFS analysed the data. AM generated the interaction networks. JFS, SMR and ESL wrote the manuscript.

