## Supplementary data for "Genetic diversity, predictive protein structures, and interaction networks of Cysteine-Rich Receptor-Like Kinases in *Arabidopsis thaliana*"

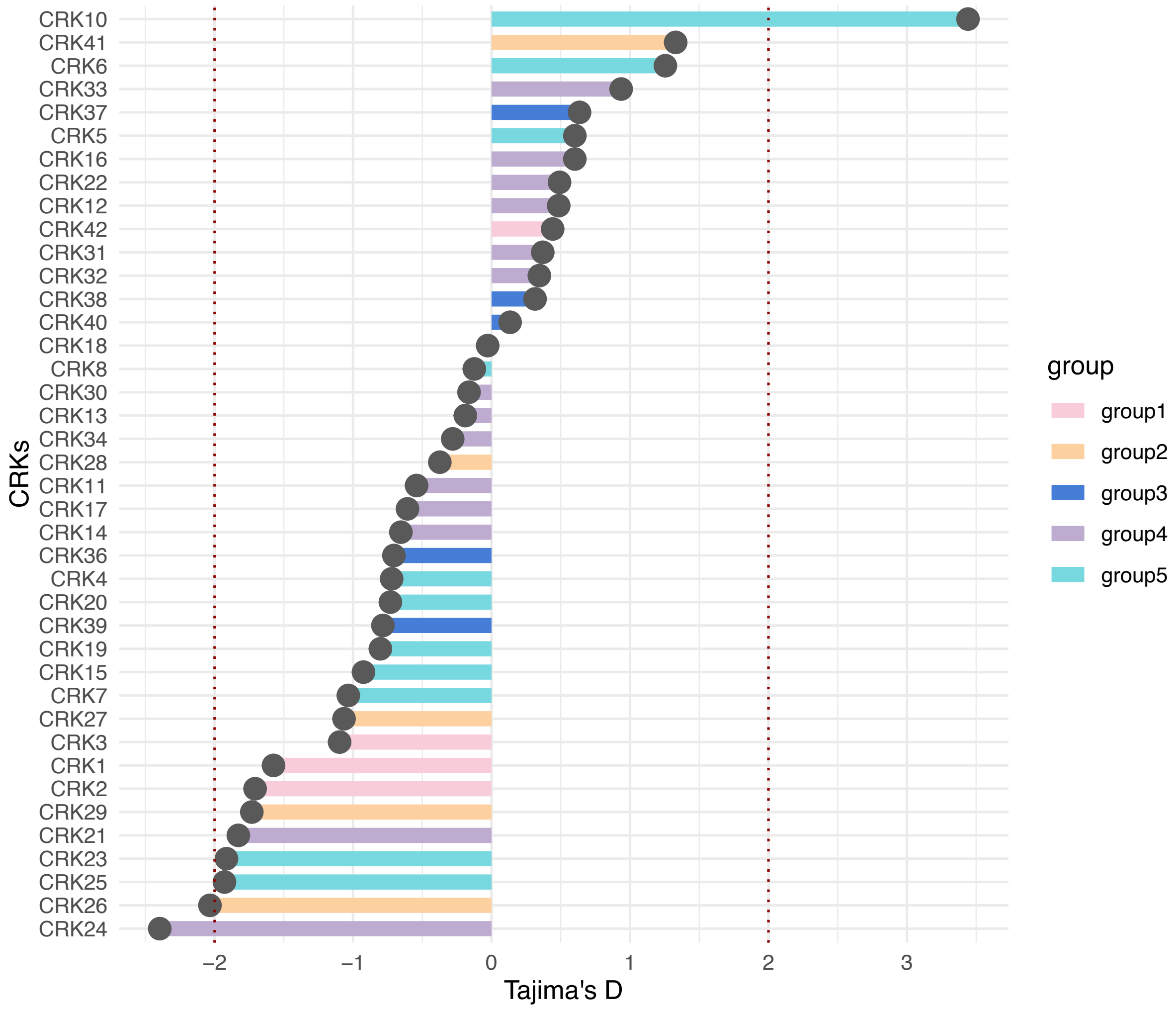


**Supplementary Figure 1.** Distribution of Tajima's D over 40 CRKs. For each CRK, the sequences were extracted across the 69 *A. thaliana* ecotypes. A dashed line was used to highlight a D value of +2 and -1. Colors correspond to the phylogenetic clades.

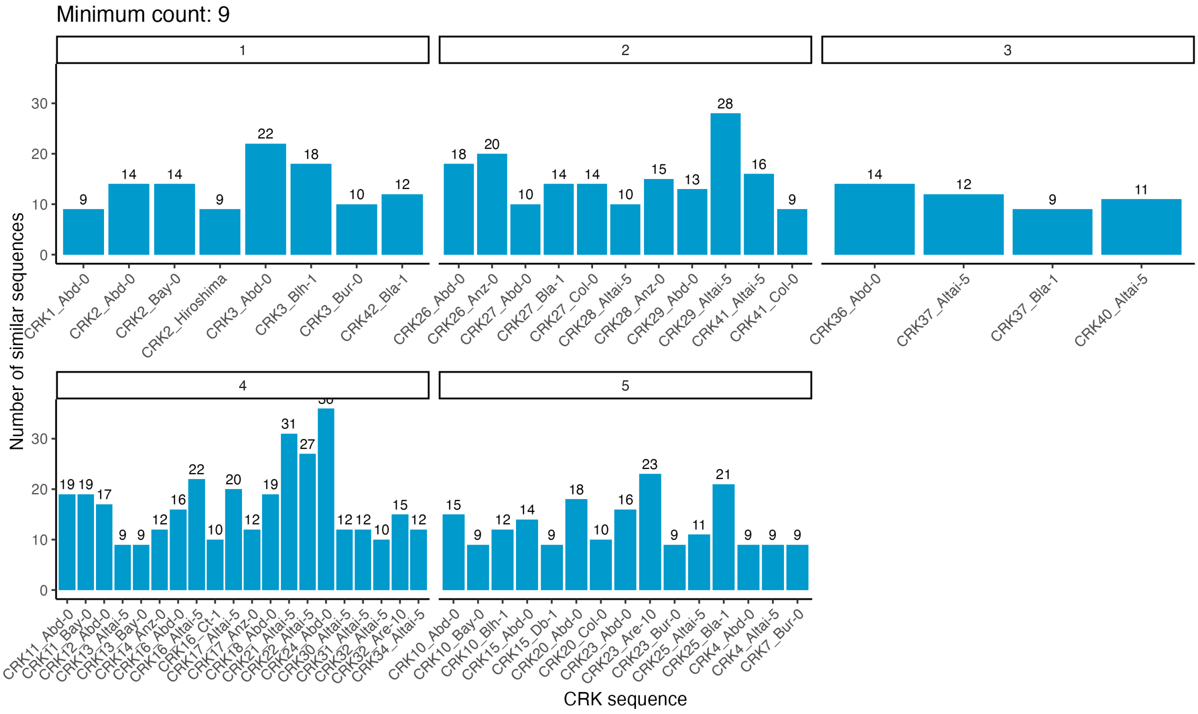


**Supplementary Figure 2.** Frequency of haplotype sequences within each phylogenetic clade. The similarity among sequences was determined in PopART with the TCS method. Similar haplotype sequences were counted (y-axis) and are represented based on a randomly picked identifier within the haplotype set (x-axis). Due to facilitating visualization, only haplotypes occurring at least nine times are displayed.

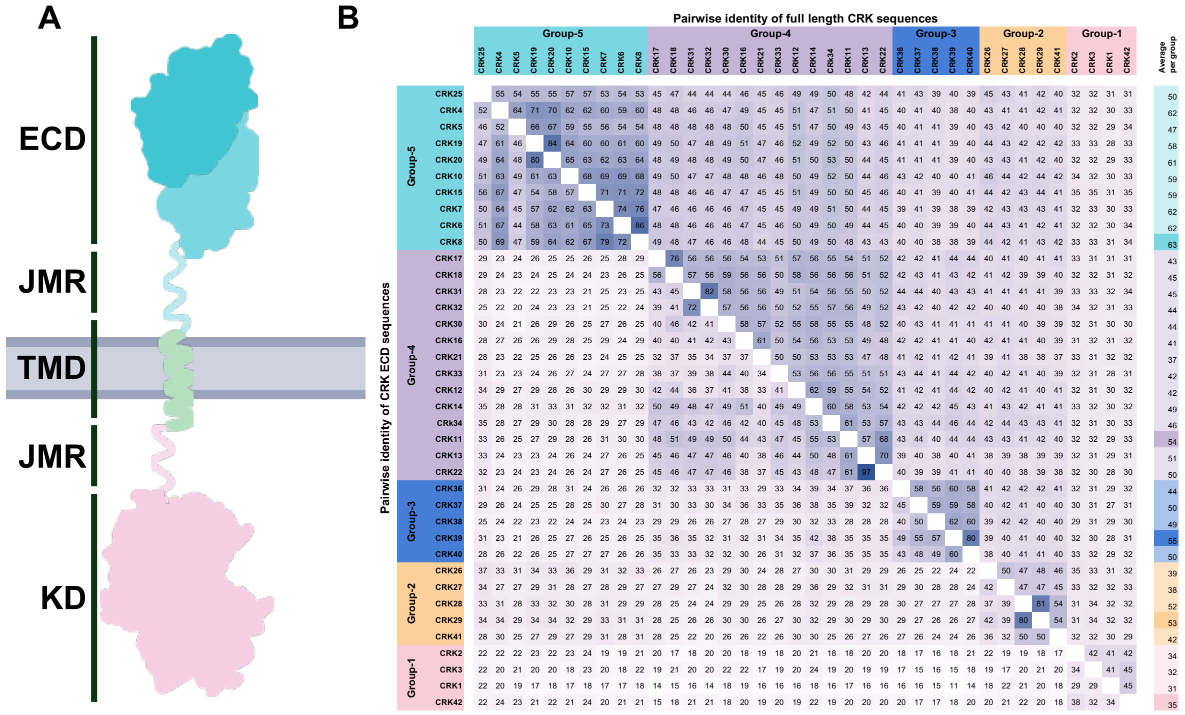


**Supplementary Figure 3. CRK-ECDs are often more sequentially diverse than the full-length protein.** **A)** Schematic of CRK structure, including the extracellular domain (ECD), transmembrane domain (TMD), the kinase domain (KD) and the two juxtamembrane regions (JMR) connecting the domains. **B)** Matrix representing the pairwise percent identity of the CRK ECDs (lower diagonal) and full-length CRKs (upper diagonal). Pairwise percent identity is the percentage of similarity of an amino acid sequence compared to the template, with high percentage meaning a high similarity. Values represent the pairwise percent identity and the squares are coloured based on the value size with high percentages (high similarity) coloured in blue and low percentages (low similarity) coloured in light pink. Pairwise identities were determined by Clustal-O (version 1.2.4). The average pairwise identity per CRK-ECD with other members of the phylogenetic group is shown in the rightmost column. The highest average pairwise identity, used to determine the representative CRKs, is marked with a darker colour.

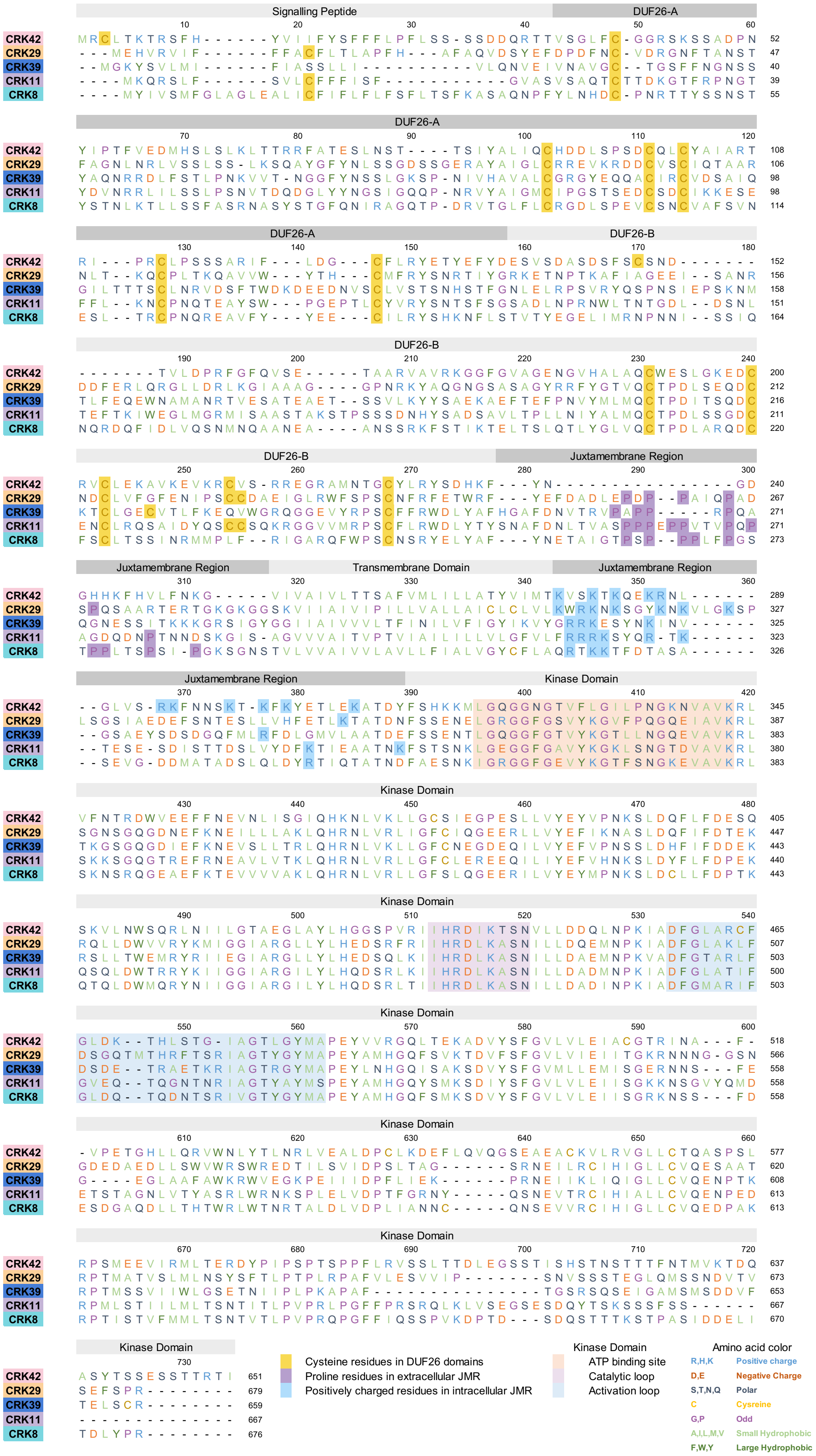

Signalling Peptide
DUF26-A
DUF26-B
Juxtamembrane Region
Transmembrane Domain
Kinase Domain
CRK42
CRK29
CRK39
CRK11
CRK8
Cysteine residues in DUF26 domains
Proline residues in extracellular JMR
Positively charged residues in intracellular JMR
Kinase Domain
ATP binding site
Catalytic loop
Activation loop
Amino acid color
R,H,K Positive charge
D,E Negative Charge
S,T,N,Q Polar
C Cysreine
G,P Odd
A,I,L,M,V Small Hydrophobic
F,W,Y Large Hydrophobic

**Supplementary Figure 4. Alignment of representative CRKs from each phylogenetic clade.** The sequence of one CRK per phylogenetic group is shown (Group-5 – CRK8, Group-4 – CRK11, Group-3 – CRK39, Group-2 – CRK29, Group-1 – CRK42). The representative CRKs were chosen based on the highest average amino acid pairwise percentage identity of each CRK-ECD to the other members of the same phylogenetic group (Supplementary Figure 3). Amino acids are coloured based on their general biophysical properties. Sequences have been assigned to specific domains or regions. Important features within a domain have been highlighted. Sequence alignment was performed with Muscle (version 3.8).

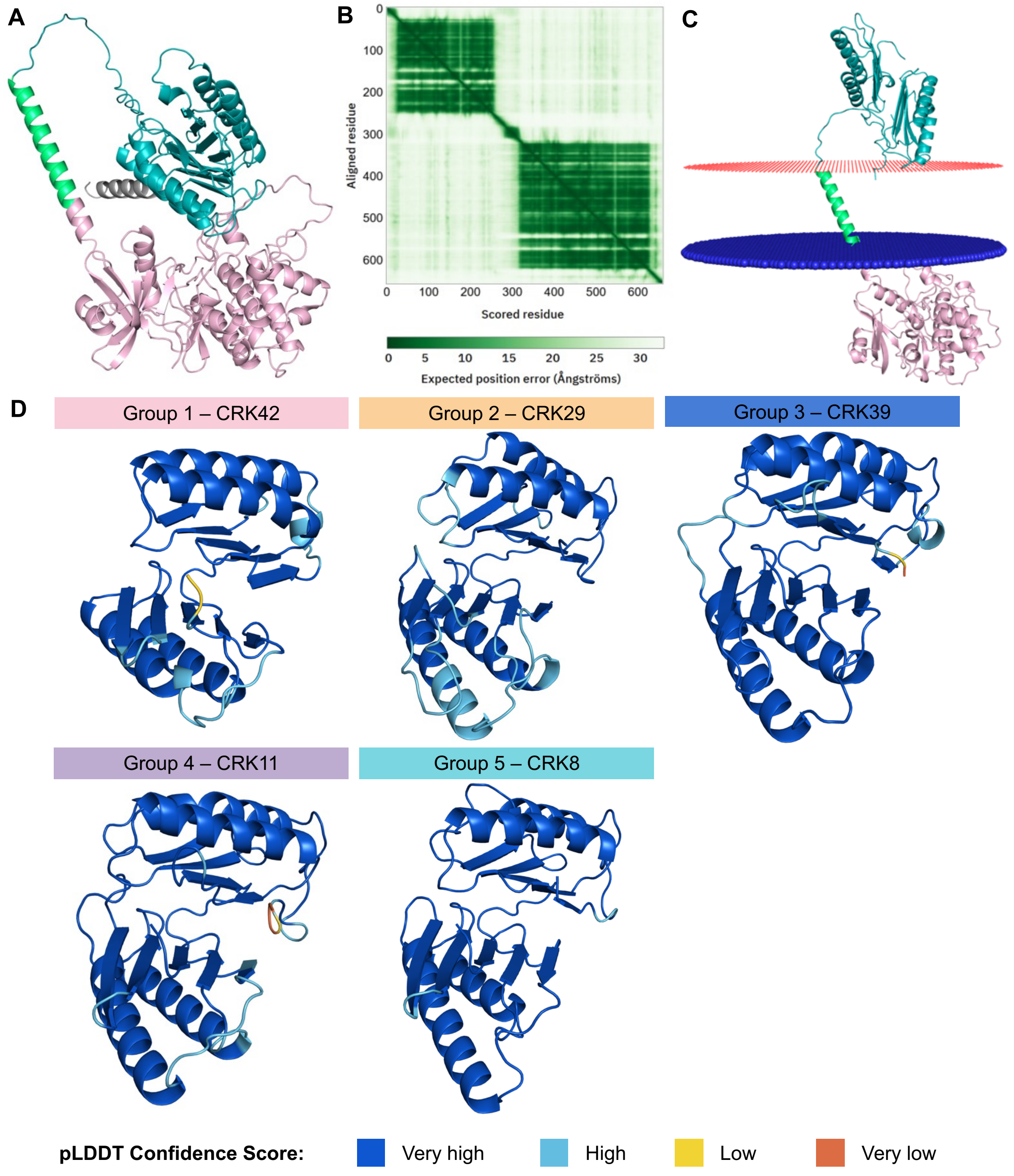


**Supplementary Figure 5. AF predicts CRK-ECDs but not the relative position of the domains towards each other with high accuracy.** A) Predicted structure of full-length CRK18 retrieved from AF database [73]. Signalling peptide indicated in grey, ECD in teal, TMD in green, and KD in pink. The ECD and KD are incorrectly placed in close proximity. B) A heatmap showing the Predicted Aligned Error (PAE) scores of CRK18. PAE scores within domains have a low error, while the PAE scores between domains are high, suggesting low confidence in the relative position of the domains. C) CRK18 model with the membrane reintroduced (in red and blue) shows separation of the ECD and the KD. The model with a membrane was obtained from Membranome [16]. D) CRK-ECD models representing each phylogenetic group. Residues are coloured by predicted local distance difference test (pLDDT) scores.

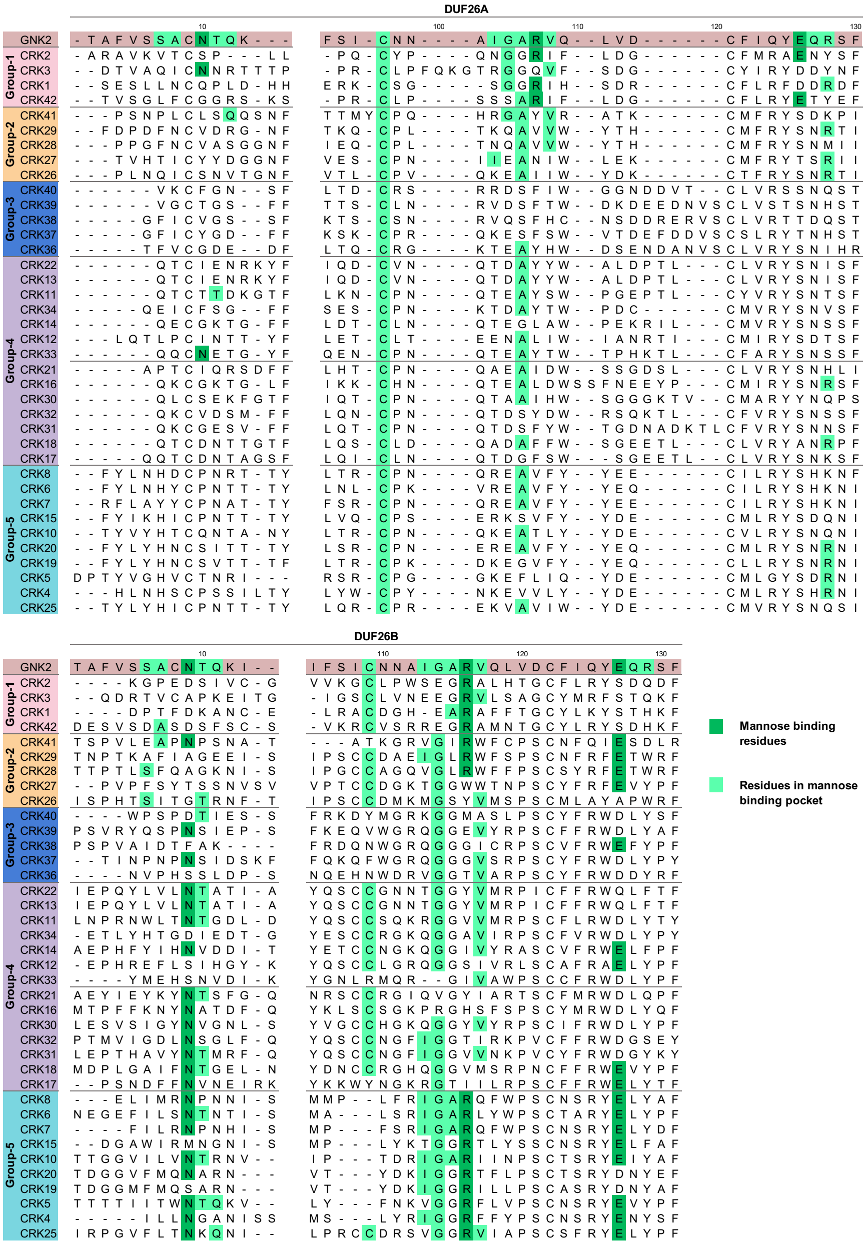


**Supplementary Figure 6. GNK2 mannose-binding residues are present in the DUF26-B domain of certain CRKs.** Sequence alignment of GNK2-mannose binding residues to CRKs DUF26-A and DUF26-B. All three mannose-binding residues are conserved in DUF26-B of Group-5 CRKs. The residues involved in GNK2-mannose binding are coloured in dark green; other residues in the binding interface are coloured in light green. Sequence alignment was performed using Muscle (version 3.8).

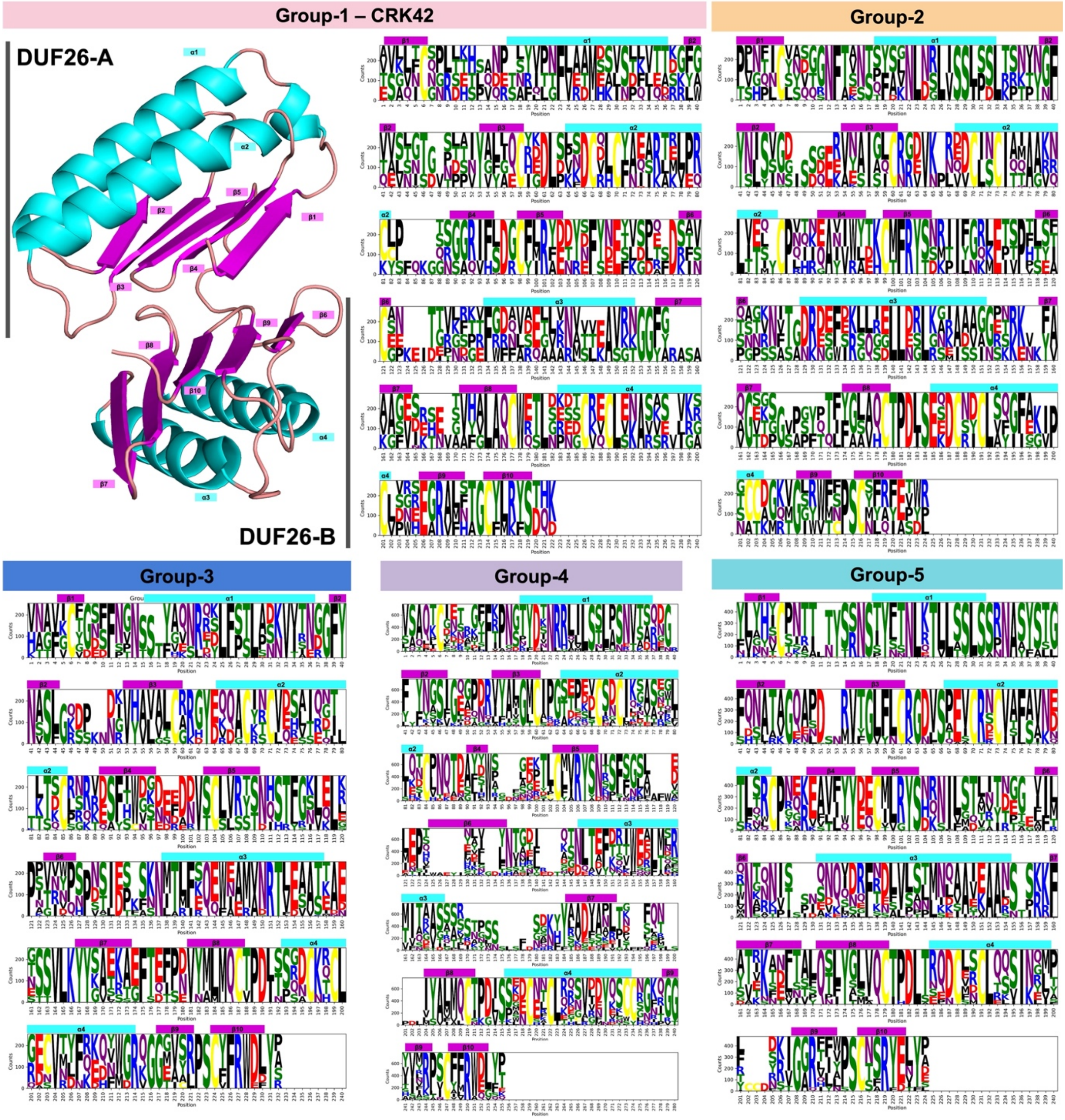


**Supplementary Figure 7. CRK Extracellular domain sequence diversity.** CRK42-ECD model consisting of two domains of unknown function 26 (DUF26-A and DUF26-B) with secondary structural elements coloured, a-helices in cyan, β-strands in magenta and loops in salmon. Logoplots of the CRK-ECD per phylogenetic group, with secondary structural elements marked (a-helices in cyan, β-strands in magenta), showing conserved and variable regions.

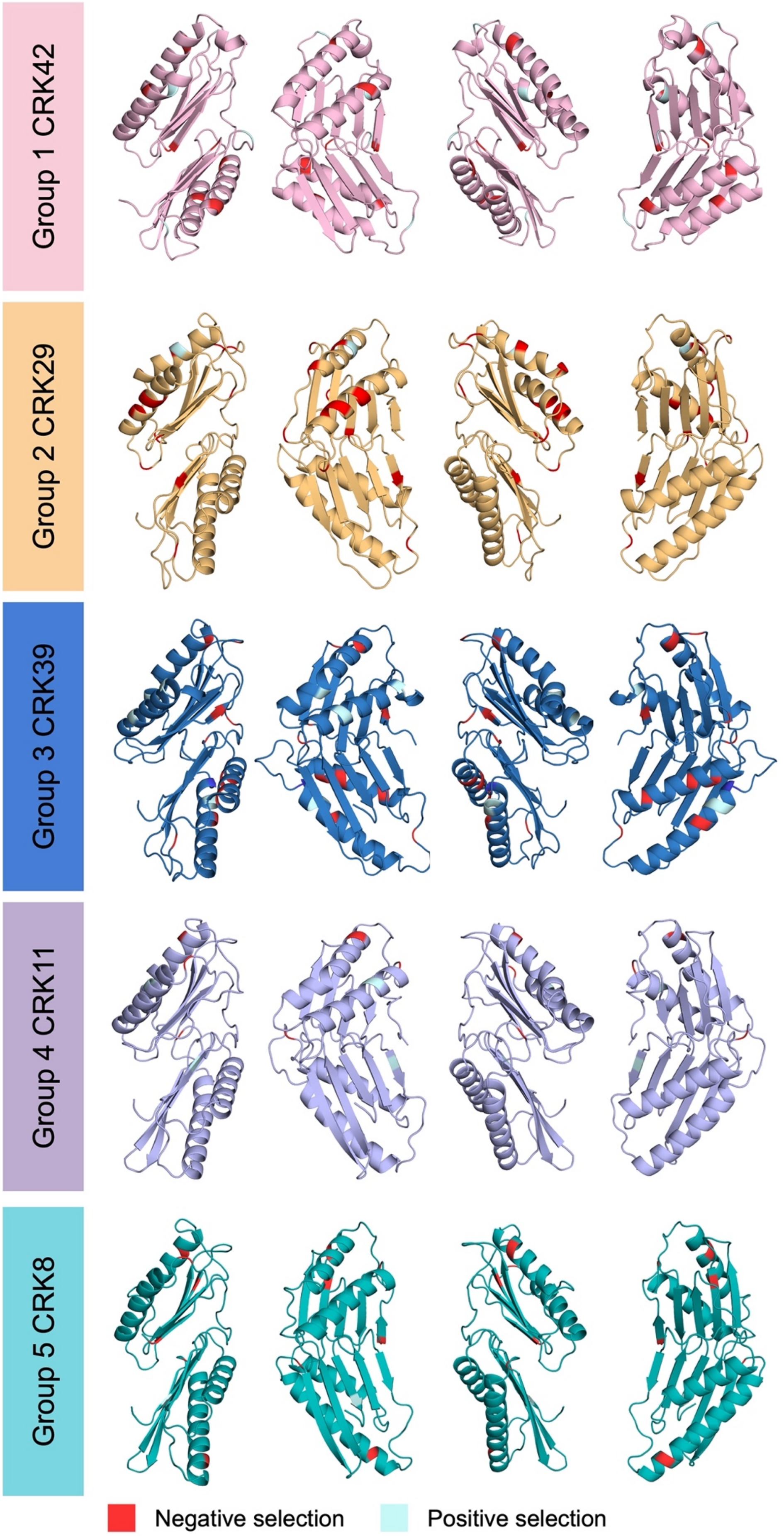


| Site | Selection |
|---|---|
| 37 | Negative |
| 56 | Negative |
| 108 | Negative |
| 168 | Negative |
| 200 | Negative |
| 213 | Negative |
| 226 | Negative |
| 81 | Positive |
| 110 | Positive |
| 143 | Positive |
| 153 | Positive |

| Site | Selection |
|---|---|
| 53 | Positive |
| 54 | Negative |
| 61 | Negative |
| 66 | Negative |
| 76 | Negative |
| 80 | Negative |
| 91 | Negative |
| 98 | Negative |
| 99 | Negative |
| 103 | Negative |
| 104 | Negative |
| 125 | Negative |
| 133 | Negative |
| 146 | Negative |
| 152 | Negative |

| Site | Selection |
|---|---|
| 30 | Negative |
| 41 | Negative |
| 70 | Negative |
| 136 | Negative |
| 152 | Negative |
| 170 | Negative |
| 209 | Negative |
| 217 | Negative |
| 220 | Negative |
| 227 | Negative |
| 53 | Positive |
| 91 | Positive |
| 102 | Positive |
| 173 | Positive |
| 174 | Positive |
| 177 | Positive |

| Site | Selection |
|---|---|
| 42 | Negative |
| 98 | Positive |
| 105 | Negative |
| 140 | Positive |
| 178 | Negative |

| Site | Selection |
|---|---|
| 40 | Negative |
| 59 | Negative |
| 91 | Negative |
| 122 | Negative |
| 168 | Negative |
| 191 | Negative |
| 250 | Positive |

**Supplementary Figure 8. Positive and negative selection of residues of representative CRK-ECDs.** Figures showing models of CRK-ECDs of representative CRKs with residues found to be under positive or negative selection coloured in blue (positive) or red (negative). Tables showing the residue position and the identified selection. The representative CRK from each phylogenetic group: CRK8 (Group 5), CRK11 (Group 4), CRK39 (Group 3), CRK29 (Group 2), and CRK42 (Group 1). The representative CRKs were chosen based on the highest average amino acid pairwise percentage identity of each CRK-ECD to the other members of the same phylogenetic group (Supplementary Figure 3).

**Supplementary Table 1**

List of CRK identifiers

| CRK | uniprot_ID | TAIR | TAIR_representative | gene_length | strand | synteny |
|---|---|---|---|---|---|---|
| CRK1 | Q9LMB9 | AT1G19090 | AT1G19090.1 | 2266 | positive | solo |
| CRK2 | Q9CAL3 | AT1G70520 | AT1G70520.1 | 2447 | negative | clusterG |
| CRK3 | Q9CAL2 | AT1G70530 | AT1G70530.1 | 2630 | negative | clusterG |
| CRK4 | Q9LZU4 | AT3G45860 | AT3G45860.1 | 2641 | negative | solo |
| CRK5 | Q9C5S8 | AT4G23130 | At4g23130.2 | 2447 | negative | solo |
| CRK6 | Q9C5S9 | AT4G23140 | At4g23140.2 | 2641 | positive | solo |
| CRK7 | Q8L7G3 | AT4G23150 | AT4G23150.1 | 2571 | positive | solo |
| CRK8 | O65468 | AT4G23160 | AT4G23160.2 | 2762 | positive | solo |
| CRK10 | Q8GYA4 | AT4G23180 | AT4G23180.1 | 2610 | positive | clusterA |
| CRK11 | Q9ZP16 | AT4G23190 | AT4G23190.1 | 2514 | negative | clusterA |
| CRK12 | O65472 | AT4G23200 | AT4G23200.1 | 2555 | negative | clusterB |
| CRK13 | Q0PW40 | AT4G23210 | AT4G23210.3 | 2527 | negative | clusterB |
| CRK14 | Q8H199 | AT4G23220 | AT4G23220.1 | 2791 | negative | clusterC |
| CRK15 | Q8W4G6 | AT4G23230 | AT4G23230.1 | 2453 | negative | clusterC |
| CRK16 | O65476 | AT4G23240 | AT4G23240.1 | 2692 | negative | clusterC |
| CRK17 | Q8L710 | AT4G23250 | AT4G23250 | 2576 | negative | clusterC |
| CRK18 | Q8RX80 | AT4G23260 | AT4G23260.1 | 2528 | negative | clusterC |
| CRK19 | Q8GWJ7 | AT4G23270 | AT4G23270 | 2913 | negative | clusterC |
| CRK20 | O65479 | AT4G23280 | AT4G23280 | 2732 | positive | clusterC |
| CRK21 | Q3E9X6 | AT4G23290 | At4g23290.2 | 2901 | negative | clusterC |
| CRK22 | Q6NQ87 | AT4G23300 | AT4G23300.1 | 2530 | positive | clusterC |
| CRK23 | O65482 | AT4G23310 | AT4G23310.1 | 3027 | positive | clusterC |
| CRK24 | O65483 | AT4G23320 | AT4G23320 | 2109 | negative | clusterC |
| CRK25 | Q9M0X5 | AT4G05200 | AT4G05200 | 2517 | negative | solo |
| CRK26 | Q9T0J1 | AT4G38830 | AT4G38830.1 | 2605 | positive | solo |
| CRK27 | O49564 | AT4G21230 | AT4G21230.1 | 2436 | negative | solo |
| CRK28 | O65405 | AT4G21400 | AT4G21400.1 | 2492 | negative | clusterD |
| CRK29 | Q8S9L6 | AT4G21410 | AT4G21410 | 2188 | negative | clusterD |

| | | | | | | |
|---|---|---|---|---|---|---|
| CRK30 | Q9LDT0 | AT4G11460 | AT4G11460.1 | 2626 | positive | clusterE |
| CRK31 | Q9LDM5 | AT4G11470 | AT4G11470 | 2433 | positive | clusterE |
| CRK32 | Q9LDS6 | AT4G11480 | AT4G11480.1 | 2392 | positive | clusterE |
| CRK33 | Q9LDN1 | AT4G11490 | AT4G11490.1 | 2701 | positive | clusterE |
| CRK34 | Q9LDQ3 | AT4G11530 | AT4G11530.1 | 2507 | positive | solo |
| CRK36 | Q9XEC6 | AT4G04490 | AT4G04490.1 | 2682 | negative | clusterF |
| CRK37 | Q9XEC7 | AT4G04500 | AT4G04500.1 | 2357 | positive | clusterF |
| CRK38 | Q9XEC8 | AT4G04510 | AT4G04510 | 2535 | positive | clusterF |
| CRK39 | Q9SYS7 | AT4G04540 | AT4G04540.1 | 2559 | positive | clusterF |
| CRK40 | Q9SYS3 | AT4G04570 | AT4G04570.1 | 2673 | positive | clusterF |
| CRK41 | O23081 | AT4G00970 | AT4G00970.1 | 3258 | positive | solo |
| CRK42 | Q9FNE1 | AT5G40380 | AT5G40380.1 | 2918 | positive | solo |

**Supplementary Table 2. Overview of analysis of Alphafold2 CRK-ECD pairs.** The left side of the table shows an analysis of the number of interactions per phylogenetic group and per CRK-ECD. Per CRK-ECD, the number (#) of interactions that pass the confidence score cut-offs was determined, and the % of the maximum number of interactions (38) was calculated. Per phylogenetic group, the total number of interactions of all CRK-ECDs in the group was determined, and the average was calculated. On the right-hand side of the table, the orientations of the interactions that passed the cut-offs were analysed. For each orientation (Standard, Flipped, Flipped and shifted, and Other), the number of CRK-ECD pairs were counted. In addition, for each phylogenetic group and CRK, the number of pairs in each orientation was counted. From the number of pairs per orientation, the percentage (%) of pairs in that orientation versus the total number of pairs per CRK or in the phylogenetic group was calculated.

### # CRK-ECDs:

| | # | % |
|---|---|---|
| Tested | 38 | |
| Pairs | 741 | |
| Pass cut-offs | 145 | 20 |

### Orientation:

| | # | % |
|---|---|---|
| Standard | 113 | 78 |
| Flipped | 21 | 14 |
| Flipped and shifted | 11 | 8 |

### Per phylogenetic group:

| Group | # CRKs | # Interactions | Average | Standard | % | Flipped | % | Flipped and shifted | % |
|---|---|---|---|---|---|---|---|---|---|
| Group-5 | 10 | 129 | 13 | 107 | 83 | 13 | 10 | 9 | 7 |
| Group-4 | 14 | 93 | 7 | 79 | 85 | 5 | 5 | 9 | 10 |
| Group-3 | 5 | 20 | 4 | 1 | 5 | 16 | 80 | 3 | 15 |
| Group-2 | 5 | 36 | 7 | 31 | 86 | 4 | 11 | 1 | 3 |
| Group-1 | 4 | 5 | 1 | 1 | 20 | 4 | 80 | 0 | 0 |

### Per CRK:

| CRK | # Interactions | % of total | Homodimer | Standard | % | Flipped | % | Flipped and shifted | % |
|---|---|---|---|---|---|---|---|---|---|
| CRK25 | 14 | 37 | 0 | 14 | 100 | 0 | 0 | 0 | 0 |
| CRK4 | 11 | 29 | 1 | 9 | 82 | 0 | 0 | 2 | 18 |
| CRK5 | 8 | 21 | 1 | 5 | 63 | 2 | 25 | 1 | 13 |
| CRK19 | 24 | 63 | 1 | 20 | 83 | 3 | 13 | 1 | 4 |
| CRK20 | 21 | 55 | 1 | 18 | 86 | 2 | 10 | 1 | 5 |
| CRK10 | 14 | 37 | 0 | 14 | 100 | 0 | 0 | 0 | 0 |
| CRK15 | 5 | 13 | 0 | 5 | 100 | 0 | 0 | 0 | 0 |
| CRK7 | 7 | 18 | 0 | 5 | 71 | 1 | 14 | 1 | 14 |
| CRK6 | 14 | 37 | 0 | 11 | 79 | 3 | 21 | 0 | 0 |
| CRK8 | 11 | 29 | 0 | 6 | 55 | 2 | 18 | 3 | 27 |
| CRK17 | 3 | 8 | 0 | 3 | 100 | 0 | 0 | 0 | 0 |
| CRK18 | 14 | 37 | 1 | 11 | 79 | 1 | 7 | 2 | 14 |
| CRK31 | 10 | 26 | 0 | 10 | 100 | 0 | 0 | 0 | 0 |
| CRK32 | 7 | 18 | 0 | 6 | 86 | 1 | 14 | 0 | 0 |
| CRK30 | 6 | 16 | 0 | 5 | 83 | 1 | 17 | 0 | 0 |
| CRK16 | 2 | 5 | 0 | 1 | 50 | 0 | 0 | 1 | 50 |
| CRK21 | 1 | 3 | 0 | 1 | 100 | 0 | 0 | 0 | 0 |
| CRK33 | 3 | 8 | 0 | 3 | 100 | 0 | 0 | 0 | 0 |
| CRK12 | 10 | 26 | 0 | 7 | 70 | 0 | 0 | 3 | 30 |
| CRK14 | 4 | 11 | 0 | 4 | 100 | 0 | 0 | 0 | 0 |
| CRK34 | 10 | 26 | 0 | 9 | 90 | 0 | 0 | 1 | 10 |
| CRK11 | 11 | 29 | 0 | 8 | 73 | 1 | 9 | 2 | 18 |
| CRK13 | 7 | 18 | 0 | 7 | 100 | 0 | 0 | 0 | 0 |
| CRK22 | 5 | 13 | 0 | 4 | 80 | 1 | 20 | 0 | 0 |
| CRK36 | 2 | 5 | 0 | 0 | 0 | 0 | 0 | 2 | 100 |
| CRK37 | 4 | 11 | 0 | 0 | 0 | 4 | 100 | 0 | 0 |
| CRK38 | 9 | 24 | 0 | 0 | 0 | 9 | 100 | 0 | 0 |
| CRK39 | 3 | 8 | 0 | 1 | 33 | 1 | 33 | 1 | 33 |
| CRK40 | 2 | 5 | 0 | 0 | 0 | 2 | 100 | 0 | 0 |
| CRK26 | 3 | 8 | 0 | 2 | 67 | 1 | 33 | 0 | 0 |
| CRK27 | 11 | 29 | 1 | 9 | 82 | 2 | 18 | 0 | 0 |
| CRK28 | 14 | 37 | 1 | 13 | 93 | 0 | 0 | 1 | 7 |
| CRK29 | 6 | 16 | 0 | 6 | 100 | 0 | 0 | 0 | 0 |
| CRK41 | 2 | 5 | 0 | 1 | 50 | 1 | 50 | 0 | 0 |
| CRK2 | 2 | 5 | 0 | 1 | 50 | 1 | 50 | 0 | 0 |
| CRK3 | 0 | 0 | 0 | 0 | 0 | 0 | 0 | 0 | 0 |
| CRK1 | 0 | 0 | 0 | 0 | 0 | 0 | 0 | 0 | 0 |
| CRK42 | 3 | 8 | 0 | 0 | 0 | 3 | 100 | 0 | 0 |

Supplementary data 1. Sequences of 40 CRK members from *Arabidopsis thaliana* Col-0.

Supplementary data 2. Sequence alignment of 2,760 CRK sequences. Sequences extracted for 40 CRK members across the 69 *Arabidopsis thaliana* ecotypes.
